# Unpouching Peracarida relationships with ultraconserved elements

**DOI:** 10.1101/2025.11.21.689751

**Authors:** Tammy Iwasa-Arai, Katrin Linse, Sónia C. S. Andrade, Gonzalo Giribet

## Abstract

Peracarida is a large group containing twelve orders of brooding crustaceans, including the large orders Amphipoda, Isopoda, Tanaidacea and Cumacea, and a series of smaller orders, some restricted to isolated habitats. The relationships of Peracarida have been disputed and no attempt has been made to use extensive taxon sampling with a modern genetic approach. Here we present a novel probe set of ultraconserved elements (UCEs) developed for peracarids to investigate higher-level relationships using newly collected Antarctic material and collection-based specimens. Concatenated and coalescent-based analyses across different levels of occupancy matrices recovered strong support for the monophyly of Peracarida and for the clade Mancoida (Isopoda + Tanaidacea + Cumacea). Thermosbaenacea was consistently resolved as the sister group to all remaining peracarids. Within Amphipoda, our results contrast with previous phylogenies by placing Corophiida and Hyperiidea as early-branching lineages. In Isopoda, Oniscidea was recovered as monophyletic and concordant with morphological hypotheses. These findings provide one of the first phylogenomic frameworks for Peracarida and demonstrate the promise of UCEs for resolving long-standing questions in malacostracan evolution.

## 1. Introduction

Crustaceans (the non-hexapod Pancrustacea) are amongst the most abundant and diverse invertebrate groups, with species inhabiting marine, freshwater and terrestrial environments. They display a variety of biological and ecological adaptations, including a larval phase (Chan & Høeg, 2015; Freire et al., 2021) or direct development (Poore, 2005); sessile (Chan & Høeg, 2015), free-living or parasitic lifestyles (Bernot et al., 2021; Iwasa-Arai & Serejo, 2018); meiofaunal size (Dahms & Qian, 2004) or gigantism (Li et al., 2021); among others. Moreover, they have economic value as an important food resource and as bioindicators (Bracken-Grissom and Wolfe, 2020).

The class Malacostraca encompasses some of the most popular crustaceans, including crabs, shrimps, lobsters and woodlice, and although several recent analyses have focused on the relationships among Eumalacostraca, Eucarida and Decapoda, many phylogenetic doubts remain within and across major lineages, especially regarding the internal relationships within the superorder Peracarida (Bracken-Grissom and Wolfe, 2020; Bernot et al., 2023). Peracarids constitute a large group of malacostracan crustaceans, with over 21,000 species inhabiting marine, freshwater, and terrestrial habitats (Spears et al., 2005), and also ranging all latitudes, with the highest diversity in the tropics, mid-latitudes and on the Antarctic shelf (Arfianti & Costello, 2020) (Fig. 1). They are defined by the presence of a marsupium, formed from thin flattened plates borne on the basalmost segments of the legs. Despite some groups presenting large sizes and being easily accessible, such as some species of Amphipoda and Isopoda, most peracarids are inconspicuous, inhabit remote and inaccessible places, and have received little taxonomic and phylogenetic attention.

**Figure 1.**
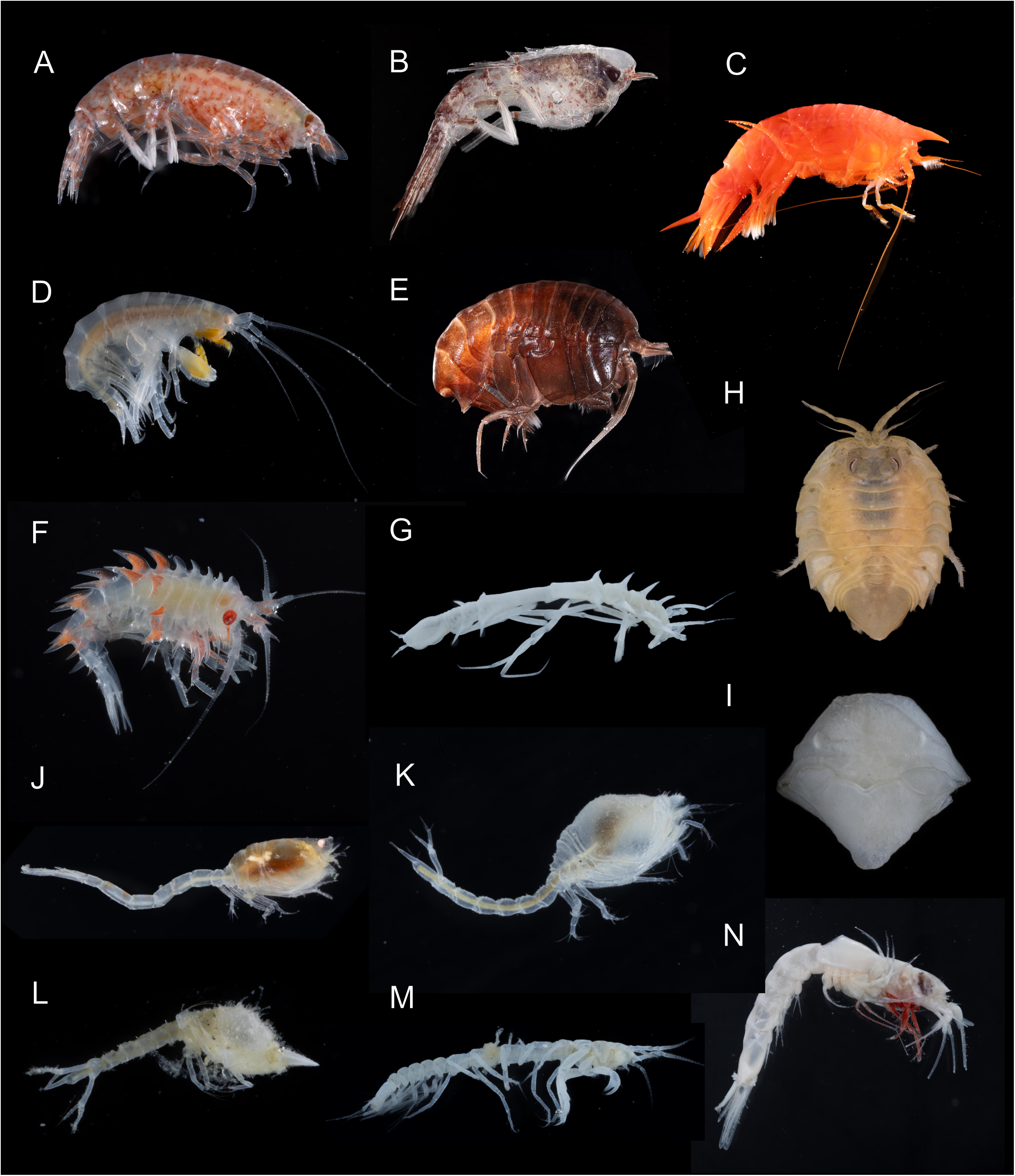
Diversity of Peracarida used in the present phylogeny. Amphipoda: **A.** *Vibilia* sp. (MCZ:IZ:172966); **B.** *Themisto gaudichaudii* (MCZ:IZ:172670); **C.** *Cyphocaris richardi* (MCZ:IZ:172829); **D.** Eusiridae sp. (MCZ:IZ:172955); **E.** *Parandania boecki* (MCZ:IZ:172673); **F.** *Acanthonotozomoides oatesi* (MCZ:IZ:172843); Isopoda: **G.** Ischnomesidae (MCZ:IZ:172929)**; H.** Serolidae sp. (MCZ:IZ:172911); **I.** Sphaeromatidae (MCZ:IZ:172980); Cumacea: **J.** Lampropidae sp. (BAS SD046-B-0276); **K.** Diastylidae (MCZ:IZ:172926); **L.** *Vemakylindrus* sp. (MCZ:IZ:172964); Tanaidacea: **M.** Apseudidae sp. ( MCZ:IZ:172928); Mysida: **N.** Erythropinae sp. (MCZ:IZ:172927).

Indeed, not only relationships between the orders Amphipoda, Bochusacea, Cumacea, Ingolfiellida, Isopoda, Lophogastrida, Mictacea, Mysida, Spelaeogriphacea, Stygiomysida, Tanaidacea and Thermosbaenacea are still disputed (e.g., Wilson, 2009; Wirkner and Richter, 2010; Robin et al., 2021), but also the interfamilial relationships of the most diverse groups, Amphipoda, Isopoda and Tanaidacea (e.g., Wilson, 2009; Drumm, 2010; Copilaș-Ciocianu et al., 2020).

The debates on the monophyly of Peracarida and the relationships between its orders have long attracted interest and controversy. Most phylogenetic proposals were published in the late 20th century, yet no consensus was reached regarding the orders’ relationships (Fig. 2; e.g., Pires, 1987; Wagner, 1994; Schram and Hof, 1998; Hessler and Watling, 1999; Richter and Scholtz, 2001). Richer and Scholtz (2001) coded 91 morphological characters to examine eumalacostracan relationships, with strong emphasis on Peracarida. Poore (2005) provided a second parsimony-based approach using 92 morphological characters, compiled *de novo*, this time focusing exclusively on Peracarida, and recovering ‘Mysidacea.’ Spears et al. (2005) proposed, in the same year, the first molecular phylogeny of Peracarida, based on 18S rRNA data, and these remained some of the most comprehensive peracarid phylogenies until now. Additional morphological characters based on the circulatory organ system were later added to the Richter and Scholtz (2001) matrix, totaling 110 morphological characters and 22 characters from the circulatory system (Wirkner and Richter, 2010). In comparison with Poore (2005), Wirkner and Richter (2010) also recovered ‘Mysidacea’, and supported the ‘Mancoida’ clade. Jenner et al. (2009) analyzed a novel morphological data matrix and molecular data in the context of eumalacostracan phylogeny, and Robin et al. (2021) added fossil taxa to Jenner et al.’s (2009) matrix. Since then, no analysis focusing specifically on peracarid relationships has been conducted. Therefore, not only relationships between the orders Amphipoda, Bochusacea, Cumacea, Ingolfiellida, Isopoda, Lophogastrida, Mictacea, Mysida, Spelaeogriphacea, Stygiomysida, Tanaidacea and Thermosbaenacea are still disputed (e.g., Wilson, 2009; Wirkner and Richter, 2010; Robin et al., 2021), but also the interfamilial relationships of the most diverse groups, Amphipoda, Isopoda and Tanaidacea (e.g., Wilson, 2009; Drumm, 2010; Copilaș-Ciocianu et al., 2020).

**Figure 2.**
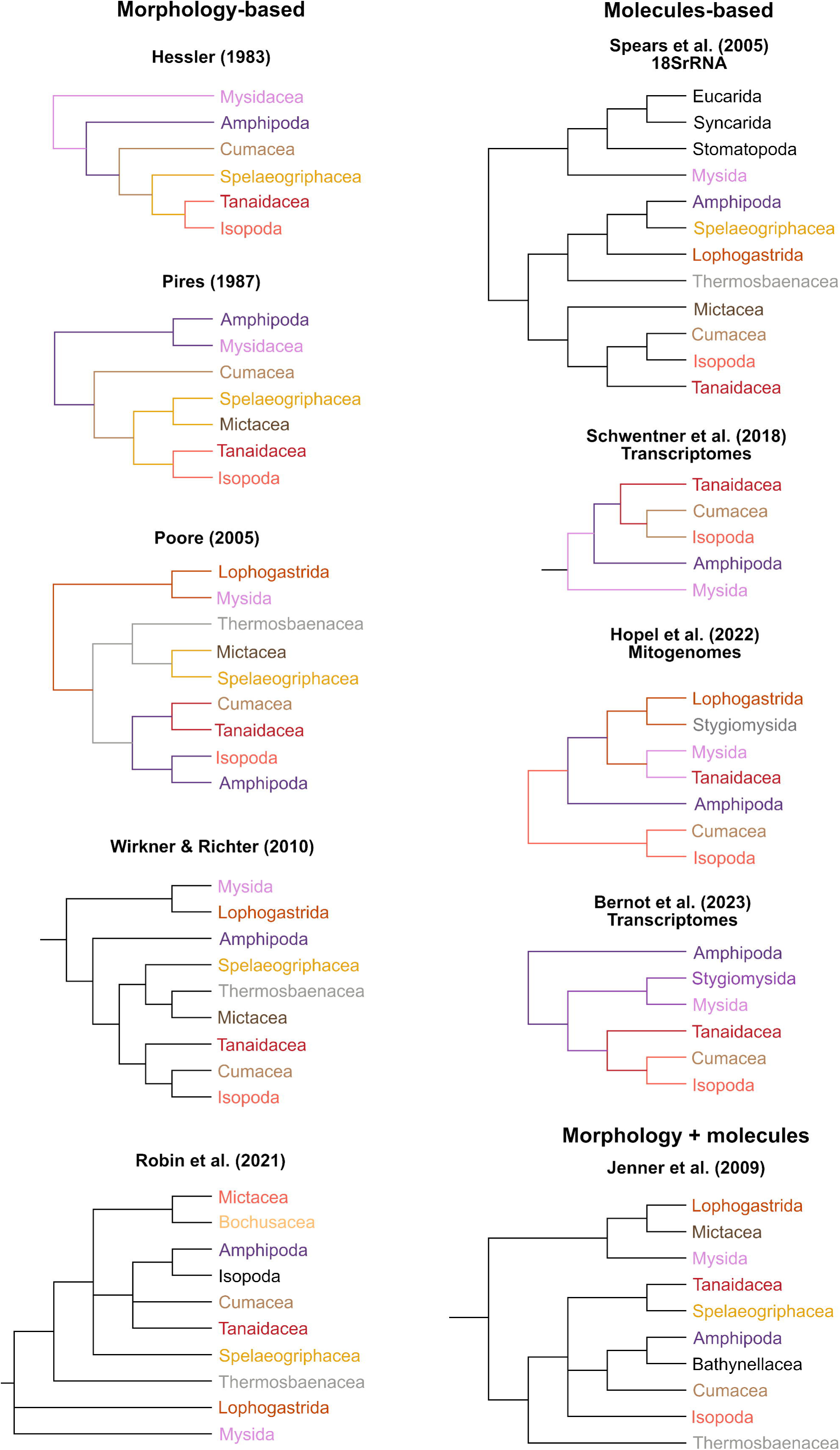
Phylogenies including Peracarida based on morphological data: Hessler (1983), Pires (1987), Poore (2005), Wirkner and Richter (2010), Robin et al. (2021); Molecular data: Spears et al. (2005), Schwentner et al. (2018), Höpel et al. (2022), Bernot et al. (2023); and Morphology and molecular data: Jenner et al. (2009).

Neither morphology nor molecules-based phylogenies have so far found a congruent and well-supported topology for Peracarida. While a few proposals have been made targeting Peracarida using morphological characters, molecular-based approaches have only been used to infer higher taxon affinities, with most attention focused towards Pancrustacea and Malacostraca (e.g. Regier et al., 2005, 2010; Schwentner et al., 2018; Höpel et al., 2022; Bernot et al., 2023), within orders (Wilson, 2009; Drumm, 2010; Copilaș-Ciocianu et al., 2020; Gerken et al., 2022) or within the families (e.g., Boyko et al., 2013; Sotka et al., 2017; Copilaș-Ciocianu et al., 2019; Iwasa-Arai et al., 2025). To date, the only phylogenomic approaches applied to understanding peracarid phylogeny are presented by Schwentner et al. (2018) and Bernot et al. (2023), but not all orders were represented. Likewise, the mitogenomic analysis of Höpel et al. (2022) lacked key taxa, like Thermosbaenacea, Mictacea, Bochusacea and Spelaeogriphacea.

While phylogenomic reconstructions based on malacostracan and decapod relationships first focused on phylotranscriptomics (Schwentner et al., 2017, 2018; Lozano-Fernandez et al., 2019), most of the orders of Peracarida remained underrepresented in phylogenomic analyses, especially by the difficulty of obtaining fresh tissues for RNA extraction. In contrast, specimens from natural history collections remain a great source of knowledge, not only for taxonomic and biogeographic studies, but also as a diversity reservoir in genomic studies, especially for rare taxa. In this context, ultraconserved elements (UCEs) are a useful tool to reconstruct phylogenies with high diversity of taxa and low cost, and they have become one of the most popular tools for invertebrate phylogenomics, from the most (Arthropoda, Faircloth et al., 2015) to the least speciose taxa (Priapulida, Raeker et al., 2025). UCEs have also been rsuccessfully used as ain phylogenomic and population studies of crustaceans. Geburzi et al. (2024) published the first UCE probe set for crustaceans, where they recovered over 1,000 loci, focusing on the main lineages within Decapoda, but they were also shown to be informative to resolve phylogenetic relationships among other malacostracans. These results indicated the usefulness of UCEs in crustaceans of difficult sampling, including several peracarid orders.

Here, we build on prior work on crustacean genomics and designed a new UCE probe set for Peracarida. By combining data from published genomes, published transcriptomes and newly collected UCEs from recent specimens, we propose a new phylogenetic hypothesis for Peracarida and show the potential for resolving the internal relationships of the large orders Isopoda and Amphipoda.

## 2. Material and Methods

### 2.1. Taxon sampling

For the probe set design, we used 20 available genomes comprising the orders Amphipoda, Isopoda, Mysida and Thermosbaenacea, with a focus on Amphipoda. The sandhopper *Morinoia aosen* (Amphipoda, Talitridae) and the skeleton shrimp *Caprella mutica* (Amphipoda, Caprellidae) were chosen as the base genome for its high level of completeness and phylogenetic interest. Table 1 shows the genomes used as exemplar taxa. Three Decapoda genomes were used as the outgroup taxon (Table 1). We downloaded 111 additional peracarid transcriptomes from NCBI to test the probe set in-silico, including the orders Amphipoda, Isopoda, Mysida, Stygiomysida and Tanaidacea (Supplementary Material Table S1). Most of the transcriptomes were secondarily removed from the analyses due to locus low retention (<10), -including representatives of Mysida (Praunus_flexuosus_SRR5140159), Stygiomysida (S_cokei_SRR5140123), and Tanaidacea (Supplementary Material Table S1).

**Table 1.** Genomes used for probe set design and test. *Morinoia aosen* GCA_030386875.1 and *Caprella mutica* GCA_947561585.1 (Amphipoda) were used as base genomes. “Peracarida”, “Malacostraca”and “Metazoa” correspond to the “raw” number of loci recovered with probe sets from the ^1^present study, ^2^Geburzi et al. 2024 and ^3^Derkarabetian et al., submitted, respectively.

| Order | Species | Suborder | Family | Code | NCBI access. no. | Peracarida<sup>1</sup> | Malacostraca<sup>2</sup> | Metazoa<sup>3</sup> |
| --- | --- | --- | --- | --- | --- | --- | --- | --- |
| Amphipoda | *Caprella mutica* | Senticaudata | Caprellidae | Cmut | GCA_947561585.1 | 260 | 0 | 3 |
| Amphipoda | *Gammarus roeselii* | Senticaudata | Gammaridae | Groe | GCA_016164225.1 | 806 | 3 | 2 |
| Amphipoda | *Morinoia aosen* | Senticaudata | Talitridae | Maos | GCA_030386875.1 | 1624 | 1 | 2 |
| Amphipoda | *Platorchestia sp.* | Senticaudata | Talitridae | Plat | GCA_014220935.1 | 1699 | 2 | 3 |
| Amphipoda | *Phronima sedentaria* | Hyperideia | Phronimidae | Psed | GCA_037179465.1 | 101 | 15 | 1 |
| Amphipoda | *Parhyale hawaiiensis* | Senticaudata | Hyalidae | Phaw | GCA_001587735.2 | 1531 | 1 | 1 |
| Amphipoda | *Trinorchestia longiramus* | Senticaudata | Talitridae | Tlon | GCA_006783055.1 | 1684 | 1 | 3 |
| Amphipoda | *Hyaella sp.* | Senticaudata | Hyaellidae | Hyal | GCA_039513755.1 | 1651 | 1 | 3 |
| Amphipoda | *Orchestia grillus* | Senticaudata | Talitridae | Ogri | GCA_014899125.1 | 1701 | 0 | 3 |
| Amphipoda | *Hyaella azteca* | Senticaudata | Hyaellidae | Hazt | GCA_000764305.4 | 1674 | 2 | 2 |
| Decapoda | *Eriocheir sinensis* | Brachyura | Varunidae | Esin | GCA_024679095.1 | 86 | 1167 | 0 |
| Decapoda | *Penaeus vannamei* | Dendrobranchiata | Penaidae | Pvan | GCA_042767895.1 | 56 | 1098 | 0 |
| Decapoda | *Callinectes sapidus* | Brachyura | Portunidae | Csap | GCA_020233015.1 | 149 | 1026 | 3 |
| Isopoda | *Armadillidium nasatum* | Oniscidea | Armadillidiidae | Anas | GCA_009176605.1 | 139 | 2 | 2 |
| Isopoda | *Trachelipus rathkii* | Oniscidea | Trachelipodidae | Trat | GCA_015478945.1 | 90 | 2 | 1 |
| Isopoda | *Porcellio scaber* | Oniscidea | Porcellionidae | Psca | GCA_034700385.1 | 100 | 2 | 0 |
| Isopoda | *Bathynomus jamesi* | Cymothoida | Cirolanidae | Bjam | GCA_023014485.1 | 32 | 5 | 0 |
| Isopoda | *Haplophthalmus danicus* | Oniscidea | Trichoniscidae | Hdan | GCA_034700045.1 | 40 | 2 | 0 |
| Isopoda | *Trichoniscus pusillus* | Oniscidea | Trichoniscidae | Tpus | GCA_034700725.1 | 36 | 1 | 0 |
| Isopoda | *Ligia exotica* | Oniscidea | Ligiidae | Lexo | GCA_002091915.1 | 28 | 4 | 0 |
| Isopoda | *Ceratothoa steindachneri* | Cymothoida | Cymothoidae | Cste | GCA_963969495.1 | 29 | 3 | 0 |
| Isopoda | *Asellus aquaticus* | Asellota | Asellidae | Aaqu | GCA_964212115.1 | 19 | 3 | 0 |
| Isopoda | *Notopais sp. 1* | Asellota | Munnopsidae | Npol | GCA_028455955.1 | 10 | 2 | 0 |
| Isopoda | *Notopais sp. 2* | Asellota | Munnopsidae | Ncry | GCA_028455975.1 | 9 | 1 | 0 |
| Mysida | *Archaeomysis grebnitzkii* |  | Mysidae | Agre | GCA_031471125.1 | 14 | 0 | 0 |

For the in-vitro test of the probe set, we compiled 48 samples from the collections of the Museum of Comparative Zoology, Cambridge, MA (MCZ), and the Smithsonian National Museum of Natural History, Washington DC (USNM), comprising seven of the peracarid orders (Table 2).

**Table 2.** Samples and description of final UCE data after assembly and filtering steps.

| Species | Order | Suborder | Family | Voucher | UC E no. | Source | Total bp | Mean length | 95 CI length | Min length | Max length | Median length | Contigs > 1kb |
| --- | --- | --- | --- | --- | --- | --- | --- | --- | --- | --- | --- | --- | --- |
| *Acanthonotozomoides oatesi* | Amphipoda | Amphilochidea | Acanthonotozomellidae | MCZ:IZ:172843 | 477 | UCE Library | 138,923 | 534.32 | 12.26 | 94 | 1,207 | 547.0 | 1121 |
| *Boeckxelia potanini* | Amphipoda | Senticaudata | Acanthogammaridae | SRR3467044 | 21 | Transcriptome | 8,897 | 741.42 | 111.32 | 215 | 1,120 | 783.5 | 5 |
| *Caprella mutica* | Amphipoda | Senticaudata | Caprellidae | GCA_947561585.1 | 241 | Genome | 14,658 | 1127.54 | 42.96 | 968 | 1,616 | 1112.0 | 11 |
| *Caprella penantis* | Amphipoda | Senticaudata | Caprellidae | USNM1463333 | 250 | UCE Library | 18,271 | 264.80 | 7.96 | 99 | 450 | 270.0 | 98 |
| *Cyamus boopis* | Amphipoda | Senticaudata | Cyamidae | MCZ:IZ:173261 | 251 | UCE Library | 209,538 | 1301.48 | 70.00 | 78 | 4,327 | 1223.0 | 19257 |
| *Cyphocaris richardi* | Amphipoda | Amphilochidea | Cyphocarididae | MCZ:IZ:172829 | 484 | UCE Library | 57,090 | 326.23 | 6.87 | 96 | 788 | 314.0 | 1665 |
| *Deutella incerta* | Amphipoda | Senticaudata | Caprellidae | USNM1605057 | 254 | UCE Library | 120,455 | 700.32 | 25.93 | 108 | 2,665 | 669.5 | 2519 |
| *Dyopedos* sp. | Amphipoda | Senticaudata | Dulichiiidae | USNM1644295 | 135 | UCE Library | 51,186 | 550.39 | 135.74 | 100 | 12,966 | 389.0 | 104 |
| *Epimeria* sp. | Amphipoda | Amphilochidea | Epimeriidae | MCZ:IZ:172924 | 183 | UCE Library | 23,165 | 335.72 | 53.64 | 87 | 3,493 | 263.0 | 304 |
| *Eulimnogammarus vittatus* | Amphipoda | Senticaudata | Eulimnogammaridae | SRR3467061 | 32 | Transcriptome | 69,281 | 769.79 | 36.22 | 201 | 1,121 | 814.0 | 38 |
| *Eurythenes obesus* | Amphipoda | Amphilochidea | Eurythenidae | MCZ:IZ:172669 | 28 | UCE Library | 14,649 | 771.00 | 384.63 | 86 | 5,681 | 253.0 | 71 |
| *Gammarus roeseli* | Amphipoda | Senticaudata | Gammaridae | GCA_016164225.1 | 714 | Genome | 16,613 | 1107.53 | 11.56 | 946 | 1,120 | 1120.0 | 14 |
| *Hyachelia tortugae* | Amphipoda | Senticaudata | Hyalidae | MCZ:IZ:173266 | 380 | UCE Library | 105,121 | 611.17 | 22.09 | 87 | 1,776 | 596.5 | 861 |
| *Hyaella azteca* | Amphipoda | Senticaudata | Hyalellidae | GCA_000764305.4 | 1463 | Genome | 1,925,374 | 1150.16 | 0.74 | 828 | 1,254 | 1160.0 | 1657 |
| *Hyaella* sp. | Amphipoda | Senticaudata | Hyalellidae | GCA_039513755.1 | 1472 | Genome | 1,569,492 | 950.63 | 6.05 | 292 | 1,185 | 1068.0 | 896 |
| *Hyperia curvicephala* | Amphipoda | Hyperidea | Hyperidae | MCZ:IZ:172965 | 75 | UCE Library | 19,849 | 336.42 | 12.92 | 96 | 658 | 312.0 | 385 |
| *Laticorophium baconi* | Amphipoda | Corophiida | Corophiidae | USNM1605092 | 148 | UCE Library | 24,160 | 284.24 | 10.75 | 87 | 603 | 264.0 | 48 |
| *Morinoia aosen* | Amphipoda | Senticaudata | Talitridae | GCA_030386875.1 | 1409 | Genome | 1,874,104 | 1154.00 | 0.91 | 731 | 1,201 | 1160.0 | 1600 |
| *Neoxenodice* sp. | Amphipoda | Senticaudata | Podoceridae | MCZ:IZ:172934 | 274 | UCE Library | 73,586 | 465.73 | 17.66 | 79 | 1,253 | 443.5 | 520 |
| *Orchestia grillus* | Amphipoda | Senticaudata | Talitridae | GCA_014899125.1 | 1482 | Genome | 1,948,393 | 1145.44 | 1.55 | 205 | 1,204 | 1160.0 | 1660 |
| *Parandania boeckii* | Amphipoda | Amphilochidea | Stegocephalidae | MCZ:IZ:172673 | 345 | UCE Library | 54,018 | 380.41 | 10.97 | 96 | 786 | 367.0 | 728 |
| *Parhyale hawaiiensis* | Amphipoda | Senticaudata | Hyalidae | GCA_001587735.2 | 1349 | Genome | 1,736,886 | 1134.48 | 2.12 | 388 | 1,414 | 1160.0 | 1456 |
| *Phronima sedentaria* | Amphipoda | Hyperidea | Phronimidae | GCA_037179465.1 | 98 | Genome | 75,778 | 750.28 | 28.15 | 249 | 1,160 | 745.0 | 27 |
| *Phronima* sp. | Amphipoda | Hyperidea | Phronimidae | MCZ:IZ:173260 | 50 | UCE Library | 5,014 | 250.70 | 31.84 | 96 | 604 | 258.0 | 362 |
| *Platorchestia* sp. | Amphipoda | Senticaudata | Talitridae | GCA_014220935.1 | 1476 | Genome | 1,954,262 | 1150.24 | 0.99 | 609 | 1,214 | 1160.0 | 1674 |
| *Podocerus brasiliensis* | Amphipoda | Senticaudata | Podoceridae | USNM1605825 | 224 | UCE Library | 25,965,037 | 249.42 | 0.78 | 78 | 7,775 | 171.0 | 2017 |
| *Themisto gaudichaudii* | Amphipoda | Hyperidea | Hyperidae | MCZ:IZ:172670 | 113 | UCE Library | 59,086 | 642.24 | 28.15 | 92 | 1,437 | 659.0 | 1097 |
| *Trinorchestia longiramus* | Amphipoda | Senticaudata | Talitridae | GCA_006783055.1 | 1684 | Genome | 1,934,670 | 1148.85 | 1.02 | 726 | 1,208 | 1160.0 | 1658 |
| *Vibilia* sp. | Amphipoda | Hyperidea | Vibiliidae | MCZ:IZ:172966 | 130 | UCE Library | 49,363 | 425.54 | 19.22 | 96 | 1,023 | 373.0 | 674 |
| Ampeliscidae sp. | Amphipoda | Amphilochidea | Ampeliscidae | MCZ:IZ:172983 | 141 | UCE Library | 20,373 | 299.60 | 10.97 | 93 | 693 | 269.5 | 1089 |
| Eusiridae sp. | Amphipoda | Amphilochidea | Eusiridae | MCZ:IZ:172955 | 465 | UCE Library | 116,962 | 512.99 | 18.23 | 89 | 2,026 | 475.5 | 3288 |
| Nannastacidae sp. | Cumacea |  | Nannastacidae | MCZ:IZ:172885 | 56 | UCE Library | 25,207 | 365.32 | 12.20 | 86 | 727 | 369.0 | 664 |
| Lampropidae sp. | Cumacea |  | Lampropidae | MCZ:IZ:172887 | 76 | UCE Library | 69,800 | 691.09 | 35.77 | 86 | 2,147 | 617.0 | 1957 |
| Leuconoide sp. | Cumacea |  | Leuconidae | MCZ:IZ:172891 | 94 | UCE Library | 75,370 | 638.73 | 48.81 | 86 | 5,445 | 590.0 | 1626 |
| Diastylidae sp. | Cumacea |  | Diastylidae | MCZ:IZ:172889 | 118 | UCE Library | 59,644 | 420.03 | 23.40 | 87 | 2,312 | 355.0 | 2011 |
| *Callinectes sapidus* | Decapoda | Brachyura | Portunidae | GCA_020233015.1 | 135 | Genome | 168,871 | 1133.36 | 6.39 | 528 | 1,173 | 1160.0 | 144 |
| *Eriocheir sinensis* | Decapoda | Brachyura | Varunidae | GCA_024679095.1 | 74 | Genome | 19,846 | 1102.56 | 13.47 | 883 | 1,122 | 1120.0 | 17 |
| *Armadillidium nasatum* | Isopoda | Oniscidea | Armadillidae | GCA_009176605.1 | 118 | Genome | 147,677 | 1062.42 | 12.26 | 496 | 1,160 | 1126.0 | 111 |
| *Bathynomus jamesi* | Isopoda | Cymothoida | Cirolanidae | GCA_023014485.1 | 28 | Genome | 35,215 | 1100.47 | 15.11 | 776 | 1,164 | 1120.0 | 30 |
| *Ceratothoa steindachneri* | Isopoda | Cymothoida | Cymothoidae | GCA_963969495.1 | 27 | Genome | 32,665 | 1126.38 | 4.56 | 1,073 | 1,164 | 1120.0 | 29 |
| *Eurycope* sp. | Isopoda | Asellota | Munnopsidae | MCZ:IZ:172910 | 105 | UCE Library | 97,453 | 792.30 | 31.29 | 86 | 1,593 | 856.0 | 2439 |
| *Glyptonotus antarcticus* | Isopoda | Valvifera | Chaetiidae | MCZ:IZ:140281 | 73 | UCE Library | 12,364,776 | 274.38 | 1.18 | 78 | 9,404 | 253.0 | 546 |
| *Haploniscus* sp. | Isopoda | Asellota | Haploniscidae | MCZ:IZ:172849 | 98 | UCE Library | 93,464 | 753.74 | 30.21 | 101 | 1,869 | 734.0 | 1013 |
| *Haplophthalmus danicus* | Isopoda | Oniscidea | Trichoniscidae | GCA_034700045.1 | 37 | Genome | 38,621 | 965.53 | 30.46 | 426 | 1,161 | 1034.0 | 23 |
| *Ianthopsis* sp. | Isopoda | Asellota | Acanthaspidiidae | MCZ:IZ:172931 | 35 | UCE Library | 4,253,563 | 260.83 | 2.01 | 78 | 5,400 | 240.0 | 289 |
| *Idotea* sp. | Isopoda | Valvifera | Idoteidae | MCZ:IZ:166902 | 130 | UCE Library | 144,368 | 995.64 | 53.50 | 98 | 4,922 | 908.0 | 11301 |
| *Ligia exotica* | Isopoda | Oniscidea | Ligiidae | GCA_002091915.1 | 25 | Genome | 2,1992 | 785.43 | 49.75 | 307 | 1,149 | 784.0 | 8 |
| *Natatolana* sp. | Isopoda | Cymothoida | Cirolanidae | MCZ:IZ:172842 | 124 | UCE Library | 72,110 | 550.46 | 20.59 | 79 | 1,578 | 516.0 | 1252 |
| *Porcellio scaber* | Isopoda | Oniscidea | Porcellio scaber | GCA_034700385.1 | 72 | Genome | 88,516 | 885.16 | 25.64 | 239 | 1,161 | 937.0 | 42 |
| *Rocinela* sp. | Isopoda | Cymothoida | Aegidae | MCZ:IZ:173268 | 86 | UCE Library | 34,354 | 373.41 | 12.06 | 96 | 681 | 349.0 | 2572 |
| *Trachelipus rathkii* | Isopoda | Oniscidea | Trachelipodidae | GCA_015478945.1 | 90 | Genome | 95,262 | 1058.47 | 14.61 | 567 | 1,163 | 1119.5 | 69 |
| *Trichoniscus pusillus* | Isopoda | Oniscidea | Trichoniscidae | GCA_034700725.1 | 34 | Genome | 2,6240 | 728.89 | 44.36 | 182 | 1,151 | 713.0 | 7 |
| *Vanhoeffenura* sp. | Isopoda | Asellota | Munnopsidae | MCZ:IZ:172827 | 95 | UCE Library | 69,937 | 647.56 | 28.59 | 94 | 2,209 | 636.5 | 541 |
| Gnathiidae sp. | Isopoda | Cymothoida | Gnathiidae | MCZ:IZ:172918 | 80 | UCE Library | 56,996 | 599.96 | 34.42 | 100 | 1,423 | 512.0 | 294 |
| Mesosignidae sp. | Isopoda | Asellota | Mesosignidae | MCZ:IZ:172923 | 86 | UCE Library | 37,820 | 393.96 | 21.16 | 100 | 1,709 | 329.5 | 148 |
| Ischnomesidae sp. | Isopoda | Asellota | Ischnomesidae | MCZ:IZ:172930 | 89 | UCE Library | 43,552 | 506.42 | 21.75 | 90 | 1,250 | 493.5 | 254 |
| Ischnomesidae sp. | Isopoda | Asellota | Ischnomesidae | MCZ:IZ:172929 | 91 | UCE Library | 34,361 | 452.12 | 26.64 | 87 | 1,283 | 386.0 | 359 |
| Munnidae sp. | Isopoda | Asellota | Munnidae | MCZ:IZ:172905 | 99 | UCE Library | 50,061 | 505.67 | 19.96 | 231 | 980 | 488.0 | 152 |
| Acanthaspidiidae sp. | Isopoda | Janiroidea | Acanthaspidiidae | MCZ:IZ:172845 | 104 | UCE Library | 25,245,028 | 783.73 | 51.55 | 4,163 | 667.5 | 15 | 1605 |
| Desmosomatidae sp. | Isopoda | Asellota | Desmosomatidae | MCZ:IZ:172908 | 106 | UCE Library | 46,627 | 475.79 | 35.76 | 91 | 3,524 | 442.0 | 255 |
| Munnopsidae sp. | Isopoda | Asellota | Munnopsidae | MCZ:IZ:172914 | 108 | UCE Library | 64,802 | 526.85 | 24.74 | 87 | 1,807 | 477.0 | 137 |
| Serolidae sp. | Isopoda | Sphaeromatidea | Serolidae | MCZ:IZ:172911 | 114 | UCE Library | 38,577 | 381.95 | 17.12 | 94 | 917 | 333.0 | 718 |
| Sphaeromatidae sp. | Isopoda | Sphaeromatidea | Serolidae | MCZ:IZ:172956 | 129 | UCE Library | 80,586 | 567.51 | 19.32 | 232 | 1,685 | 514.0 | 1601 |
| *Nebalia longicornis* | Leptostraca | Nebaliacea | Nebaliidae | MCZ:IZ:172977 | 91 | UCE Library | 31,004 | 378.10 | 18.55 | 86 | 879 | 356.0 | 493 |
| *Mictocaris halope* | Mictacea |  | Mictocarididae | MCZ:IZ:135174 | 84 | UCE Library | 34,717 | 399.05 | 16.25 | 230 | 1,308 | 372.0 | 329 |
| *Antarctomysis maxima* | Mysida |  | Mysidae | MCZ:IZ:140283 | 87 | UCE Library | 29,629 | 375.05 | 23.58 | 93 | 1,424 | 324.0 | 512 |
| *Archaeomysis grebnitzkii* | Mysida |  | Mysidae | GCA_031471125.1 | 12 | Genome | 89,154 | 841.08 | 27.55 | 236 | 1,121 | 907.0 | 48 |
| Mysidae sp. | Mysida |  | Mysidae | MCZ:IZ:172927 | 97 | UCE Library | 20,200 | 288.57 | 11.20 | 86 | 650 | 271.5 | 67 |
| *Zeuxo* sp. | Tanaidacea | Tanaidomorpha | Tanaididae | MCZ:IZ:173265 | 63 | UCE Library | 54,417 | 632.76 | 33.38 | 230 | 2,063 | 617.5 | 157 |
| Apseudidae sp. | Tanaidacea |  | Apseudidae | MCZ:IZ:17292 | 91 | UCE Library | 24,257,041 | 303.19 | 0.96 | 78 | 9802 | 261.0 | 1205 |
| Thermosbaenacea sp. | Thermosbaenacea |  |  | MCZ:IZ:134138 | 58 | UCE Library | 27,987 | 363.47 | 19.50 | 86 | 1,507 | 324.0 | 762 |

### 2.2. Probe set design and synthesis

We used PHYLUCE version 1.7.3 (Faircloth, 2016) to design the probe set, following the tutorial on https://phyluce.readthedocs.io/en/latest/tutorials/tutorial-4.html (also see Faircloth, 2017). The downloaded genomes were converted to FASTA format and file headers were modified for compatibility with the PHYLUCE pipeline using SeqIO, available on Bioconda (Grüning et al., 2018). Copies of the genomes in 2bit format required further downstream in the pipeline, which were created using faToTwoBit (Kent, 2002). ART (Huang et al., 2012) was then used to simulate 100-bp short reads with 200-bp insert size (150 standard deviation) from all exemplar genomes. These simulated reads were individually aligned to the base genome using minimap2 (Li, 2018), identifying putatively conserved loci with less than 5% sequence divergence to the base genome. Unmapped reads were removed from the alignment files using the view function in SAMtools (Li et al., 2009), and the cleaned alignments were converted to BED format, sorted by position along scaffolds, and proximate contigs were merged using BEDTools (Quinlan and Hall, 2010). Finally, phyluce_probe_strip_masked_loci_from_set in PHYLUCE was used to filter out putatively conserved loci that are repetitive regions, by removing all contigs shorter than 80 bp and where more than 25% of the base genome were masked.

Based on these pairwise alignment files, an SQLite database was created to query conserved loci shared across several taxa. From this database, we identified loci shared between the base genome and 15 out of 20 exemplar genomes (using *M. aosen* as the base genome) and 5 out of 20 genomes (using *C. mutica* as the base genome), and extracted sequences corresponding to these loci from the base genome, buffered to 160 bp, with phyluce_probe_get_genome_sequences_from_bed. For each of these sequences, two 120-bp probes with 40 bp overlap were designed to achieve 3X tiling density at the center of the conserved locus. Potentially problematic probes with more than 25% masked bases, below 30% or above 70% GC content, as well as potential duplicate probes with more than 50% coverage and identity were subsequently removed, to create a temporary probe set.

To design the final probe set, the temporary probes were aligned back to all exemplar genomes, now also including the Decapoda outgroups, with a minimum identity of 50%. Sequences of 180 bp length were extracted from the targeted conserved loci in all taxa and written to FASTA files. Another SQLite database with the matches of the temporary probes to conserved loci across all taxa was created and used to identify loci that were recovered by a temporary probe in at least 15 out of the 23 taxa (20 exemplar, 3 outgroup, 1 base genome using *M. aosen*). For each of these loci, two 120-bp probes with 3X tiling density were designed from all taxa, filtered and duplicates removed as described above. In-silico tests of the UCE probe set were performed by assessing locus recovery across the genomes, using phyluce_assembly_match_contigs_to_probes after aligning the probe set to these genomes (see Table 1). Subsequently, after the step of phyluce_assembly_match_contigs_to_probes, we merged the probe sets and filtered for probe duplicates. We kept more unique (shared with less taxa) probes from the *C. mutica* probe set (5 out of 20 genomes) because we were initially interested in Corophiida relationships. We then used the Decapoda probe set (Geburzi et al., 2024) for filtering probes that captured Amphipoda UCEs from reads using BOWTIE2 (Langmead and Salzberg, 2012) to add to our final probe set, building a chimera probe set based on *M. aosen*, *C. mutica* and *Eriocheir sinensis*.

The concatenated probe set file was sent to Arbor BioSciences, where final tests of the design were performed before synthesizing the probes. Each probe was BLASTed against the base genome, and hybridization was simulated under standard myBaits ® (Arbor BioSciences, MI, USA) conditions, to identify and remove non-specific probes, as well as probes targeting over-represented regions.

### 2.3. DNA extraction, library preparation and UCE sequence capture

Genomic DNA was extracted from dissected pereopods in large-bodied specimens (>10 mm) or from the whole body in small-bodied specimens. DNA extraction followed the protocol of Tin et al. (2014) using silica-based magnetic beads, with in-lab modifications (Derkarabetian et al., 2019), and a final elution volume of 50 μL in ddH_2_O. All extractions were quantified on a Qubit 2.0 fluorometer using a dsDNA High Sensitivity kit (Life Technologies, Inc.).

Libraries were prepared using the KAPA HyperPlus kit, following the manufacturer’s protocol with some in-lab modifications (see Derkarabetian et al., 2019; Moles and Giribet, 2021; Geburzi et al., 2024), particularly for the historical samples. We used up to 250 ng of DNA as input material; however, input from historical samples was usually much lower (down to 4 ng). Fresh samples were enzymatically fragmented to a target length of 500–700 bp, using 5 μL KAPA Fragmentation Enzyme, 2.5 μL KAPA Fragmentation Buffer (10X) and 17.5 μL DNA sample for a final volume of 25 μL, with incubation times between 3 and 8 min at 37°C. Older historical samples did not require fragmentation as they were naturally degraded, and 25 μL of the eluted DNA went directly into end-repair and A-tailing. End-repair and A-tailing was conducted using the KAPA HyperPlus enzyme mix for fresh, enzymatically fragmented samples, and KAPA HyperPrep enzyme mix for historical, naturally fragmented samples, with 30 min incubation at 65°C. This step was immediately followed by adapter ligation, using 10 μM universal iTru stubs and 45 min incubation at 20°C for fresh, high-input samples, and 5 μM stubs and up to 60 min incubation at 20°C for historical and low-input samples. A post-ligation cleanup was carried out using freshly prepared Serapure SpeedBeads (Rohland and Reich, 2012) with 0.8X beads for fresh, and up to 3X beads for old and low-input samples. Fifteen μL of ligated libraries were used in library amplification, with 25 μL 2X KAPA HiFi HotStart ReadyMix, 5 μL individual i5/i7 dual indexing adaptors (Glenn et al., 2019), and the following thermal protocol: 45 s at 98°C, 20 cycles of 15 s at 98°C, 30 s at 60°C, 60 s at 72°C, and 5 min at 72°C final extension. Amplified libraries were purified with SpeedBeads (1X for high-, 3X for low-input samples), quantified, and pooled into equimolar batches of eight samples with 250 ng DNA per sample. If necessary, pools were speed-vacuumed to a final volume of 14 μL.

Hybridization followed the myBaits ® Hybridization Capture for Targeted NGS manual v 5.01 with the following modifications: Hybridization time for pools was 4 h at 62°C, 16 h at 60°C and 4 h at 55°C. Fifteen μL of hybridized pools were amplified for 20 cycles using the same thermal protocol as described above, purified with AMPure beads (1.8X for pools of high-, 3X for pools of low-input samples), quantified on a Qubit 2.0, and size estimated on an Agilent TapeStation 2200. A final 1X bead cleanup was performed on pools where adapter-dimers were present. Amplified, hybridized pools were pooled in equimolar amounts and sequenced on an Illumina NovaSeq platform (paired-end, 150 bp) at the Bauer Core Facility, Faculty of Arts and Sciences, Harvard University.

### 2.4. Bioinformatics and phylogenetic analyses

Raw Illumina reads were demultiplexed and processed using PHYLUCE version 1.7.2 following the workflow in the online tutorial (https://phyluce.readthedocs.io/en/latest/tutorials/tutorial-1.html). Adapters and low-quality bases were removed with Illumiprocessor (Faircloth, 2013), which implements Trimmomatic (Bolger et al., 2014). Contigs were assembled with SPAdes v3.15.5 (Prjibelski et al., 2020) using the “–careful” option to reduce the number of mismatches and indels. Probes were matched to the assembled contigs with phyluce_assembly_match_contigs_to_probes, with a 65% threshold value for minimum locus coverage and identity. UCE sequences from all taxa were extracted to individual FASTA files per locus, including incomplete loci that were recovered only in a subset of the taxa. At this step, we also included contigs from the genomes and transcriptomes used for probe set design and in-silico tests, “harvested” with phyluce_probe_slice_sequence_from_genomes (see Table 2 for UCE recovery summary statistics for these samples).

Because of the disparity in locus occupancy among UCE library samples, genomes, and transcriptomes, most of the transcriptomes were secondarily removed to increase locus retention, and only two amphipod transcriptomes were used in the final dataset, to increase taxon sampling (Table 2), totaling 71 terminal taxa. Raw reads were trimmed and cleaned using Seqyclean v1.10.09, removing adapter sequences, vectors, oligonucleotides and reads with average Phred (QS) quality below 24 and smaller than 50 bp. Trinity v2.1.1 (Grabherr et al., 2011) was used for assembly with a pair distance set to 800. Quality statistics were calculated using BUSCO v5.4.7 (Manni et al., 2021) and the TrinityStats.pl script available on Trinity.

Extracted sequences were aligned with MAFFT (Katoh and Standley, 2013), and alignments were trimmed with GBlocks (Castresana, 2000; Talavera and Castresana, 2007), using conservative settings, i.e. –b1 0.5 –b2 0.85 –b3 4 –b4 8, suitable for phylogenetic analyses at high taxonomic levels. We built >35%, >50% and >75% occupancy matrices (M35, M50 and M75, respectively), meaning that a UCE was included if present in at least 35%, 50% and 75% of taxa, respectively.

Phylogenies were estimated using the concatenated M35, M50 and M75 occupancy matrices in IQ-TREE v2.2.0 (Minh et al., 2020a), using UCE locus partitioning (Chernomor et al., 2016) with ModelFinder (-m MFP+MERGE), and an ultrafast bootstrap (Hoang et al., 2018) with 1500 replicates. Gene and site concordance factors (gCF and sCF) were calculated in IQ-TREE 2 (Minh et al., 2020) using the -t option in order to quantify the proportion of gene trees and informative sites supporting each internal branch and to assess topological concordance and conflict across loci.

We also implemented coalescent-based tree estimation using M35, M50 and M75 matrices in ASTRAL 5.7.5 (Zhang et al. 2018). We followed Zhang et al. (2018) and pre-processed gene trees by collapsing internal branches with UFBS <50% prior to species-tree inference with ASTRAL. Collapsed gene trees were concatenated and input into ASTRAL (Zhang et al., 2018) to produce summary species trees. The resulting trees were visualized with iTOL version 6.7.4 (Letunic and Bork, 2021), and edited in Inkscape version 1.2.

## 3. Results

### 3.1. UCE probe set *in silico* tests

Initially, the probe set after filtering and final testing included 22,479 probes targeting 1,701 loci (Supplementary Material Table S1). From these, 7,648 probes were synthesized, prioritizing those that captured the largest number of peracarid loci, along with additional probes from the Decapoda probe set that also successfully captured Amphipoda loci *in vitro*.

When comparing results *in silico* from the new Peracarida probe set with probe sets targeting Decapoda (Geburzi et al., 2024) and Metazoa (Raeker et al., 2025; Derkarabetian et al., 2025), our probe set resulted in up to 500 times more recovered loci for amphipod genomes, and up to 60 times more loci for isopod genomes (Table 1).

### 3.2. UCE recovery and *in vitro* probe set efficacy

Because of the disparity of locus occupancy among UCE library samples, genomes and transcriptomes, most of the transcriptomes were secondarily removed in order to increase locus retention, and only two amphipod transcriptomes were used in the final dataset to increase taxon sampling (Table 2).

Mean raw UCE locus recovery per sample was 292 across all samples (range 14–1,701), and mean locus length across all samples was 662 bp (range of mean per sample 249–1,301 bp). Outgroup locus recovery from Decapoda *in silico* samples ranged from 86–149, whereas the in vitro sample of *Nebalia longicornis* (Leptostraca) recovered 91 loci. Within Peracarida, UCE recovery was highest among Amphipoda, which was used as the base genome and for probe set design, followed by Isopoda. The lowest locus recovery was from the genome of *Archaeomysis grebnitzkii* SRR23998316 (n=14), but sufficient to cluster the species within the Mysida clade (Fig. 2). The trimmed and revised M35 of the Peracarida dataset contained 148 UCE loci and the concatenated alignment had a length of 24,151 bp and contained 11,506 informative sites. The M50 contained 118 UCE loci (13,932 bp, 6,705 informative sites) and the M75 contained 12 UCE loci (1,996 bp, 901 informative sites).

### 3.3. UCE phylogeny of Peracarida

Analyses of the three datasets (M35, M50 and M75) under maximum likelihood produced overall congruent topologies, and our results are thus discussed based on the concatenated-based M50 analysis from here on, as it is a good compromise of minimizing missing data with the highest taxon sampling (Fig. 3). Peracarida was supported as monophyletic in all concatenated-based analyses, with bootstrap support (UFBS hereafter) of 95%, 73% and 97% in the M35, M50 and M75 occupancy datasets respectively, although with moderate gene concordance factors (gCF) (Fig. 3, Supplementary Material Figs. S1-S2). Across the phylogeny, gCFs were generally low to moderate for deep nodes (≈30%), indicating substantial gene-tree discordance among loci, whereas shallow nodes showed higher gCF values (>70%), reflecting greater agreement among genes in recent divergences. Site concordance factors (sCF) were more evenly distributed throughout the tree, ranging from 30–60%, suggesting variable but moderate site-level support across the dataset (Supplementary Material Figs. S2-S3).

**Figure 3.**
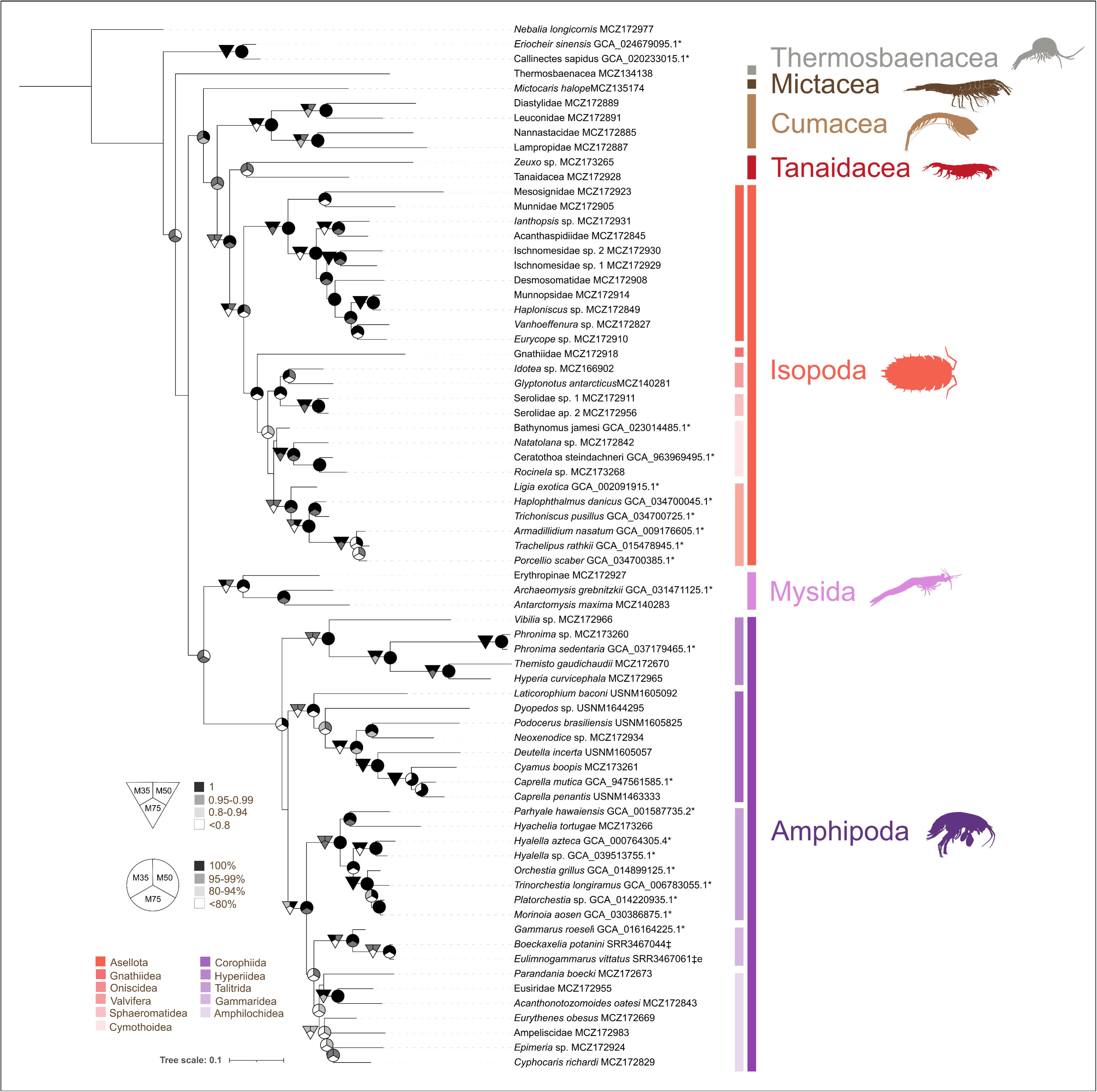
Maximum likelihood phylogeny of Peracarida inferred from M50 (50% occupancy matrix) in IQ-TREE showing ultrafast bootstrap nodal support on branches. Circles show nodes which received maximal support in concatenated-based analyses for matrices M35, M50 and M75 in IQ-TREE. Triangles show nodes with maximal support in coalescent-based analyses for matrices M35, M50 and M75 in ASTRAL. Color legends on bottom left show suborders of Amphipoda and Isopoda.

In contrast, the coalescent-based approach resulted in different relationships between orders, with M50 not recovering Peracarida, including Decapoda (Eriocheir_sinensis_GCA_024679095.1 and Callinectes_sapidus_GCA_020233015.1), but without local posterior probability (LPP) support (Supplementary Material Fig. S4).

In all analyses, of the seven peracarid sampled orders, five had multiple species represented, and were recovered as monophyletic (Fig. 3), including Amphipoda and Isopoda – the most speciose orders. Speleogriphacea, Stygiomysida and Lophogastrida were removed from the analyses because of the very low locus coverage (Supplementary Material Fig. S1), and further analyses will be needed in order to elucidate their position within Peracarida.

Major clades and order-level relationships were concordant in the three concatenated datasets (Fig. 3; Supplementary Material Fig. S1 and S2): Thermosbaenacea was found as the sister group to all other Peracarida, the latter comprising two clades: one with Mictacea, Cumacea, Tanaidacea and Isopoda; and another including Mysida and Amphipoda (Fig. 3). Mictacea was recovered as the sister clade of Cumacea+Tanaidacea+Isopoda (Mancoida), with Tanaidacea closely related to Isopoda, both these nodes supported with 100% UFBS, at least in the M50 matrix (Fig. 3).

Subordinal relationships, in contrast, differed across datasets. Within Isopoda, our analyses detected two main clades: the first, separating Asellota from other Isopoda suborders; and the second, showing a close relationship between Gnathiidea, Oniscidea, Cymothoidea, Valvifera and Sphaeromatidea (Fig. 3). We found the suborder relationships of Valvifera + Sphaeromatidea in all datasets, but this clade’s sister group varied across datasets. In all trees, we found the monophyly of the terrestrial suborder Oniscidea (Fig. 3, Supplementary Material Figs. S1-S5).

Within Amphipoda, we found Hyperiidea as the sister clade of all other amphipods in the concatenated-based M50 datasets and coalescent-based M50 and M75 datasets, with Corophiida being found either as sister clade to Hyperiidea or as the sister clade to all other amphipod suborders (Fig. 3 and Supplementary Material Figs. S3 and S5). In the concatenated-based M35, M50 and coalescent-based M50 datasets, Amphilochidea was recovered as the sister clade of Gammaridea, and Talitrida as the sister clade of Amphilochidea + Gammaridea (Fig. 3). The M75 datasets, instead, recovered Amphilochidea as a paraphyletic group, as it included the sampled gammaridean species (Supplementary Material Figs. S2 and S5). Talitrida, on the other hand, was recovered as monophyletic in all analyses (Fig. 3, Supplementary Material Figs. S1-S5). Despite these incongruences, our data suggest that Amphilochidea, Gammaridea and Talitrida are closely related amphipod clades.

## 4. Discussion

Peracarid monophyly and ingroup relationships have remained hot topics in crustacean phylogenetics for decades (Fig. 2), yet little consensus exists and few efforts have been made to investigate their relationships using genomic data, among other reasons due to the difficulties of obtaining reliable samples. Peracarids are defined by well-supported morphological characters such as the presence of a lacinia mobilis in the mandible and oostegites (flattened plates arising from the inner proximal margin of the coxa of certain pereopods) forming a marsupium, one of the main changes that allowed the most specialized adaptation to land in crustaceans (Broly et al., 2012).

The inferred relationships among the peracarid orders differ markedly across studies based on morphology, molecules, or combined data, with no order-level arrangement receiving broad agreement over time (Fig. 2). Studies based on morphological characters place Amphipoda as sister group to Mysida in Pires (1987), as the sister group to Isopoda by Poore (2005; see also Jenner et al., 2009 and Robin et al., 2021), as sister group to a clade containing Cumacea, Tanaidacea, Isopoda, Mictacea and Spelaeogriphacea in Richter and Scholtz (2001), or as sister group to a clade containing these five orders plus Thermosbaenacea in Wirkner & Richter (2010)–all based on morphological characters. Molecular phylogenies also differ between each other: Amphipoda was found to be the sister clade of Spelaeogriphacea in the 18S rRNA-based phylogeny of Spears et al. (2005), the sister clade to ((Lophogastrida + Stygiomysida) + (Mysida + Tanaidacea)) in the mitogenomic analysis of Höpel et al. (2022), and as the sister group to the remaining Peracarida in Bernot et al. (2023) maximum likelihood transcriptomic phylogeny, although Bayesian CAT-GTR analysis of the Dayhoff6 matrix in Bernot et al. (2023) recovered Amphipoda as the sister group of Mysida. In our analyses, Amphipoda was the sister group to Mysida with high UFBS support (95–99%) in all concatenated-based analyses and M35 and M75 of coalescent-based analyses, whereas Amphipoda resulted as the sister group to all other Peracarida (and Decapoda) in the M50 ASTRAL analysis, albeit without significant support (Fig. 3, Supplementary Material Figs. S1-S5).

The Mancoida clade (Isopoda + Tanaidacea + Cumacea) has higher agreement among phylogenies, and it was recovered by the molecular-based studies using 18 rRNA and transcriptomes (Spears et al. 2005; Schwentner et al., 2018; Bernot et al. 2023). Mancoida was also found in our analyses with Isopoda as the sister group to Tanaidacea (98%, 100% and 85% UFBS in the concatenated-based M75, M50 and M35 matrices, respectively), with Isopoda as the sister group to Tanaidacea likewise identified in the morphological analyses of Hessler (1983) and Pires (1987), but not with Wirkner & Richter (2010), who found Cumacea to be the sister group of Isopoda. Our results contrast with the relationships proposed by Schwentner et al. (2018: figure 2) based on phylotranscriptomics, or with the results of Spears et al. (2005), Höpel et al. (2022) and Bernot et al. (2023), that observed a closer relationship between Cumacea and Isopoda. Thermosbaenacea, one of the most singular peracarid orders including 34 known species mostly from anchialine environments (Jaume, 2008), was recovered as the sister clade to all other Peracarida, and traits such as using the dorsal part of the carapace to brood embryos suggest a deeper divergence from the remaining orders (Wagner, 1994; Olesen et al., 2015).

Of the largely sampled orders Amphipoda and Isopoda, relationships appear well supported in the M50 dataset, but do not agree with previous phylogenetic hypotheses. Copilaș-Ciocianu et al. (2020) presented the most comprehensive amphipod phylogeny to date using up to four Sanger-based markers, and found nine lineages corresponding to Gammaridea, Lysianassoidea, Crangonyctidea, Corophiidira, Eusiroidea + Iphimediidae, Physosomata + Physocephalata, Talitrida and Atylidae, with a clade comprising Gammaridea, Lysianassoidea, Crangonyctidea and Corophiidira, and another clustering Eusiroidea + Iphimediidae, Physosomata + Physocephalata and Talitrida (Copilaș-Ciocianu et al., 2020). In contrast to the two major clades found by Copilaș-Ciocianu et al. (2020), we did not find a group composed of Gammaridea and Corophiida, instead, Corophiida appears as one of the earliest divergences within Amphipoda. Moreover, we found Gammaridea (a mainly freshwater group), more closely related to Amphilochidea and Talitrida (marine, freshwater and terrestrial species), than to Corophiida, showing that environmental shifts (marine-freshwater-land) may be more prone to occur in this group, whereas Corophiida is a generally marine group with other adaptations, especially to biotic substrata, being found on sea turtles (Iwasa-Arai et al., 2020), whales (Iwasa-Arai & Serejo, 2018), sponges (Guerra-García, 2001) and algae (Duffy, 1990). Obviously, taxon sampling is missing some key amphipod families, but our relatively large sampling suggest the placement of Corophiida and Hyperiida as suborders and a closer relationship of Amphilochidea to groups within Senticaudata (Talitrida and Gammaridea).

Within Isopoda, Oniscidea has always been a group of interest, especially because of their adaptations to terrestrial environments. Recent phylogenetic work on Oniscidea suggested that the suborder is not monophyletic, and that conquest of land occurred more than once (Lins et al., 2017; Dimitriou et al., 2019). Dimitriou et al. (2019) found species of Ligiidae, a supralittoral family, as the sister clade of the remaining Oniscidea and Sphaeromatidae + Valvifera, in contrast to our results, that support monophyly of Oniscidea and suggest that Ligiidae might be the sister clade to all other oniscideans, but still part of the same lineage that gave rise to the terrestrial isopods. In contrast to the broad range of phylogenetic studies on Oniscidea and groups with peculiar ecological and evolutionary histories, such as the parasitic Epicaridea, the highly speciose suborder Asellota, and the morphologically conspicuous Serolidae (Held, 2000; Wägele et al., 2003; Raupach et al., 2009; Boyko et al., 2013) the remaining isopod orders are yet to be studied, especially in a phylogenomic context. In a broader context, Brusca and Wilson (1991) proposed a phylogeny of Isopoda based on a cladistic analysis of 92 morphological characters, encompassing the majority of isopod families and suborders, and suggested Phreatoicidea as the sister clade to all other isopods, followed by Asellota + Microcerberida, Oniscidea, and a large clade composed of the remaining families. Our results, despite the lack of some isopod families, are in overall agreement with that topology based on morphological characters, illustrating the informativeness of morphological data in phylogenetic reconstructions for the order.

The most comprehensive studies on the phylogeny of Cumacea and Tanaidacea were those of Gerken et al. (2022) and Drumm (2010), respectively, and familial relationships of cumaceans found in our concatenated-based and coalescent-based analyses disagree with the topology of Gerken et al. (2022). Of the four families analysed herein, we found a closer relationship between Diastylidae and Leuconidae, and between Lampropidae and Nannastacidae, with maximal support in all matrices, while the four locus analysis of Gerken et al. (2022) supports one clade comprising Nannastacidae and Leuconidae and one with Diastylidae and Lampropidae. Tanaidacea has been subjected to few phylogenetic studies, and our analyses only included two species and failed to provide a well-supported result. This may be due in part to the lack of tanaidacean genomes in the probe set design which could explain the low locus recovery. Peracarid phylogenetics has been a field of interest that has received little attention due to the difficulties in collecting several lineages that inhabit unique ecosystems or are found in remote locations. By leveraging a recent collection of Antarctic samples and other samples housed at the Museum of Comparative Zoology and at the Smithsonian National Museum of Natural History, we were able to develop and test a novel UCE probe set that generated dozens of genes for phylogenomic analyses. Although some orders remain unsampled (Bochusacea, Ingolfiellida, Lophogastrida, Spelaeogriphacea, Stygiomysida), our results identified high support for some novel relationships within Peracarida and show promise for future internal work in the larger orders Amphipoda and Isopoda.

## Supporting information

Table S1. Genomes, transcriptomes and UCE libraries used in the present study before locus filtering.

## Acknowledgements

Tom Illife provided samples of Mictacea and Thermosbaenacea. Adam Baldinger of the MCZ assisted with specimen work, Arianna Lord, Ella Frigyik and Ligia Benavides provided laboratory and bioinformatics training and support. Karen Osborn and Mark Lehtonen provided DNA samples from the USNM collection. Salvatore Siciliano (GEMM-Lagos) provided the *Cyamus boopis* sample. Gary Poore (Museum Victoria) identified Ischnomesidae samples. Antarctic samples from this study were obtained during the BIOPOLE-2 Cruise in February and March, 2025, and we are indebted to the crew of the RRS Sir David Attenborough and many members of the cruise who assisted with sampling. We are especially indebted to cruise leaders Sophie Fielding and Geraint Tarling and to UK Natural Environment Research Council funding for the BIOPOLE National Capability Multicentre Round 2 grant no. NE/W004933/1. We are thankful to FAPESP grants no. 2021/06738-8 and 2024/06934-0. We are indebted to one anonymous reviewer and to James Bernot for their detailed comments that helped improve this work.

## Data availability

Raw sequences are deposited in the NCBI sequence read archive (SRA) under the BioProject accession number PRJNA1333392. Assemblies, input and output files, concatenated matrices and tree files generated during the analyses are deposited on the Harvard Dataverse under https://doi.org/10.7910/DVN/AFZSHN.

## CRediT authorship contribution statement

Tammy Iwasa-Arai: Writing – original draft, Methodology, Investigation, Formal analysis. Katrin Linse: Writing – review & editing, Methodology. Sónia Andrade: Writing – review & editing, Investigation, Conceptualization. Gonzalo Giribet: Writing – review & editing, Investigation, Funding acquisition, Conceptualization.

## Supplementary Material

**Table S1.** Genomes, transcriptomes and UCE libraries used in the present study before locus filtering.

**Figure S1.** Maximum likelihood tree inferred from partitioned analysis of concatenated M35 dataset in IQTREE.

**Figure S2.** Maximum likelihood tree inferred from partitioned analysis of concatenated M75 dataset in IQTREE.

**Figure S3.** Coalescent-based tree inferred from M35 dataset in ASTRAL.

**Figure S4.** Coalescent-based tree inferred from M50 dataset in ASTRAL.

**Figure S5.** Coalescent-based tree inferred from M75 dataset in ASTRAL.

