## Supplementary material for "Unpouching Peracarida relationships with ultraconserved elements": Table S1. Genomes, transcriptomes and UCE libraries used in the present study before locus filtering.

**Table S1**. Genomes, transcriptomes and UCE libraries used in the present study before loci filtering.

| Dataset ALL | Order | Species | Voucher | raw_UCE_pera V1 | Source |
|---|---|---|---|---|---|
| Cmut | Amphipoda | *Caprella mutica* | GCA_947561585.1 | 260 | Genome |
| Groe | Amphipoda | *Gammarus roeseli* | GCA_016164225.1 | 806 | Genome |
| Esin | Decapoda | *Eriocheir sinensis* | GCA_024679095.1 | 86 | Genome |
| Csap | Decapoda | *Callinectes sapidus* | GCA_020233015.1 | 149 | Genome |
| Pvan | Decapoda | *Penaeus vannamei* | GCA_042767895.1 | 56 | Genome |
| Anas | Isopoda | *Armadillidium nasatum* | GCA_009176605.1 | 139 | Genome |
| Psed | Amphipoda | *Phronima sedentaria* | GCA_037179465.1 | 101 | Genome |
| Trat | Isopoda | *Trachelipus rathkii* | GCA_015478945.1 | 90 | Genome |
| Psca | Isopoda | *Porcellio scaber* | GCA_034700385.1 | 100 | Genome |
| Maos | Amphipoda | *Morinoia aosen* | GCA_030386875.1 | 1624 | Genome |
| Plat | Amphipoda | *Platorchestia* sp. | GCA_014220935.1 | 1699 | Genome |
| Tlon | Amphipoda | *Trinorchestia longiramus* | GCA_006783055.1 | 1684 | Genome |
| Phaw | Amphipoda | *Parhyale hawaiensis* | GCA_001587735.2 | 1531 | Genome |
| Ogri | Amphipoda | *Orchestia grillus* | GCA_014899125.1 | 1701 | Genome |
| Hyal | Amphipoda | *Hyalella* sp. | GCA_039513755.1 | 1651 | Genome |
| Hazt | Amphipoda | *Hyalella azteca* | GCA_000764305.4 | 1674 | Genome |
| Bjam | Isopoda | *Bathynomus jamesi* | GCA_023014485.1 | 32 | Genome |
| Hdan | Isopoda | *Haplophthalmus danicus* | GCA_034700045.1 | 40 | Genome |
| Tpus | Isopoda | *Trichoniscus pusillus* | GCA_034700725.1 | 36 | Genome |
| Lexo | Isopoda | *Ligia exotica* | GCA_002091915.1 | 28 | Genome |
| Cste | Isopoda | *Ceratothoa steindachneri* | GCA_963969495.1 | 29 | Genome |
| Aaqu | Isopoda | *Asellus aquaticus* | GCA_964212115.1 | 19 | Genome |
| Agre | Mysida | *Archaeomysis grebnitzkii* | SRR23998316 | 14 | Genome |
| Caprella_sp | Amphipoda | *Caprella* sp. | SRR14135878 | 93 | Transcripto |

| | | | | | |
|---|---|---|---|---|---|
| | | | | | me |
| C_dilatata | Amphipoda | *Caprella dilatata* | | 58 | Transcripto me |
| G_minus | Amphipoda | *Gammarus minus* | SRR5576336 | 172 | Transcripto me |
| M_wohli | Amphipoda | *Micruropus wahlii* | SRR3467083 | 164 | Transcripto me |
| O_albinus | Amphipoda | *Ommatogammarus albinus* | SRR3467085 | 161 | Transcripto me |
| E_similis | Amphipoda | *Eulimnogammarus similis* | SRR3467053 | 156 | Transcripto me |
| E_viridulus | Amphipoda | *Eulimnogammarus viridulus* | SRR3467060 | 157 | Transcripto me |
| A_flavus | Amphipoda | *Oxyacanthus flavus* | SRR3467038 | 152 | Transcripto me |
| Hyperia_sp | Amphipoda | *Hyperia* sp. | SRR20015048 | 152 | Transcripto me |
| E_cruentus | Amphipoda | *Eulimnogammarus cruentus* | SRR3467063 | 153 | Transcripto me |
| E_sophianosi | Amphipoda | *Heterogammarus sophianosii* | SRR3467056 | 143 | Transcripto me |
| E_czerskii | Amphipoda | *Eulimnogammarus czerskii* | SRR3467064 | 142 | Transcripto me |
| G_chevreuxi | Amphipoda | *Gammarus chevreuxi* | SRR5109803 | 138 | Transcripto me |
| G_locusta | Amphipoda | *Gammarus locusta* | SRR10868690 | 142 | Transcripto me |
| O_flavus | Amphipoda | *Oxyacanthus flavus* | SRR3467038 | 138 | Transcripto me |
| P_grubei | Amphipoda | *Pallasea grubei* | SRR3467094 | 137 | Transcripto me |
| H_setosa | Amphipoda | *Hyalellopsis setosa* | SRR3467075 | 135 | Transcripto me |
| H_grisea | Amphipoda | *Hyallelopsis grisea* | SRR3467074 | 131 | Transcripto me |
| P_cancellus | Amphipoda | *Pallasea cancellus* | SRR3467093 | 125 | Transcripto me |
| A_sowinskii | Amphipoda | *Oxyacanthus sowinskii* | SRR3467039 | 125 | Transcripto me |
| E_ussolzewi | Amphipoda | *Eulimnogammarus ussolzewii* | SRR3467054 | 123 | Transcripto me |
| M_parvulus | Amphipoda | *Micruropus parvulus* | SRR3467081 | 120 | Transcripto me |
| A_carpenteri | Amphipoda | *Boeckaxelia carpenterii* | SRR3467043 | 119 | Transcripto me |
| E_violaceus | Amphipoda | *Eulimnogammarus violaceus* | SRR3467055 | 118 | Transcripto me |
| A_grewingki | Amphipoda | *Brachyuropus grewingkii* | SRR3467040 | 116 | Transcripto me |
| P_borowskii | Amphipoda | *Parapallasea borowskii* | SRR3467098 | 117 | Transcripto me |
| B_latissima | Amphipoda | *Brandtia latissima* | SRR3467046 | 113 | Transcripto me |

| | | | | | |
|---|---|---|---|---|---|
| G_fasciatus | Amphipoda | *Gmelinoides fasciatus* | SRR3467071 | 110 | Transcriptome |
| E_vittatus | Amphipoda | *Eulimnogammarus vittatus* | SRR3467061 | 112 | Transcriptome |
| P_kessleri | Amphipoda | *Pallaseopsis kessleri* | SRR3467095 | 109 | Transcriptome |
| A_maximus | Amphipoda | *Cornugammarus maximus* | SRR3467042 | 106 | Transcriptome |
| E_wagii | Amphipoda | *Eucarinogammarus wagii* | SRR3467051 | 99 | Transcriptome |
| C_bicarinatus | Amphipoda | *Carinurus bicarinatus* | SRR3467048 | 98 | Transcriptome |
| C_inflatus | Amphipoda | *Crypturopus inflatus* | SRR3467049 | 95 | Transcriptome |
| E_veneris | Amphipoda | *Echinogammarus veneris* | SRR1050726 | 95 | Transcriptome |
| A_potanini | Amphipoda | *Boeckaxelia potanini* | SRR3467044 | 92 | Transcriptome |
| M_branickii | Amphipoda | *Macrohectopus branickii* | SRR3467077 | 88 | Transcriptome |
| P_brandtii | Amphipoda | *Homalogammarus brandtii* | SRR3467089 | 87 | Transcriptome |
| S_parasiticus | Amphipoda | *Dorogostaiskia parasitica* | SRR3467102 | 76 | Transcriptome |
| P_cancelloides | Amphipoda | *Pallasea cancelloides* | SRR3467091 | 74 | Transcriptome |
| P_podoceroides | Amphipoda | *Pandorites podoceroides* | SRR3467097 | 72 | Transcriptome |
| P_branchialis | Amphipoda | *Pachyschesis branchialis* | SRR3467088 | 69 | Transcriptome |
| E_kietlinskii | Amphipoda | *Sluginella kietlinskii* | SRR3467052 | 57 | Transcriptome |
| P_pruinosus | Isopoda | *Porcellionides pruinosus* | SRR15808558 | 49 | Transcriptome |
| N_hrabei | Amphipoda | *Niphargus hrabei* | SRR13297211 | 48 | Transcriptome |
| A_gigantea | Amphipoda | *Alicella gigantea* | SRR14866259 | 44 | Transcriptome |
| P_plebs | Amphipoda | *Pseudorchomene plebs* | SRR19909456 | 35 | Transcriptome |
| H_quarta | Amphipoda | *Halice quarta* | SRR10768829 | 35 | Transcriptome |
| B_schellenbergi | Amphipoda | *Scopelocheirus schellenbergi* | SRR14866419 | 37 | Transcriptome |
| G_japonica | Amphipoda | *Grandidierella japonica* | DRR128008 | 24 | Transcriptome |
| A_aquaticus | Isopoda | *Asellus aquaticus* | SRR13297205 | 21 | Transcriptome |
| S_terebrans | Isopoda | *Sphaeroma terebrans* | SRR4436643 | 20 | Transcriptome |
| L_hargeri | Isopoda | *Lirceus hargeri* | SRR11966484 | 14 | Transcriptome |

| | | | | | |
|---|---|---|---|---|---|
| L_culveri | Isopoda | *Lirceus culveri* | SRR11966488 | 13 | Transcriptome |
| P_cavaticus | Isopoda | *Proasellus cavaticus* | ERR3245482 | 11 | Transcriptome |
| I_baltica | Isopoda | *Idotea balthica* | SRR5140127 | 9 | Transcriptome |
| L_tripunctata | Isopoda | *Limnoria tripunctata* | SRR7059832 | 11 | Transcriptome |
| L_usdagalun | Isopoda | *Lirceus usdagalun* | SRR11966487 | 11 | Transcriptome |
| P_strouhali | Isopoda | *Proasellus strouhali* | ERR3245553 | 10 | Transcriptome |
| L_exotica | Isopoda | *Ligia exotica* | SRR14289477 | 9 | Transcriptome |
| A_hilgendorfii | Isopoda | *Asellus hilgendorfii* | SRR14135880 | 8 | Transcriptome |
| P_hercegovinensis | Isopoda | *Proasellus strouhali* | ERR1437546 | 7 | Transcriptome |
| C_spinigena | Tanaidacea | *Carpoapseudes spinigena* | SRR14135876 | 6 | Transcriptome |
| Apseudes_sp | Tanaidacea | *Apseudes* sp. | SRR14135879 | 6 | Transcriptome |
| P_algicola | Tanaidacea | *Parapseudes algicola* | SRR14135873 | 6 | Transcriptome |
| P_aragonensis | Isopoda | *Proasellus aragonensis* | ERR1433129 | 6 | Transcriptome |
| P_karamani | Isopoda | *Proasellus karamani* | ERR1437545 | 6 | Transcriptome |
| Ceratothoa_sp | Isopoda | *Ceratothoa* sp. | SRR15808556 | 5 | Transcriptome |
| G_antarcticus | Isopoda | *Glyptonotus antarcticus* | SRR4017484 | 5 | Transcriptome |
| P_grafi | Isopoda | *Proasellus grafi* | ERR1437552 | 6 | Transcriptome |
| P_ibericus | Isopoda | *Proasellus ibericus* | ERR1437535 | 10 | Transcriptome |
| P_spelaeus | Isopoda | *Proasellus spelaeus* | ERR1433130 | 4 | Transcriptome |
| P_tomiokaensis | Tanaidacea | *Phoxokalliapseudes tomiokaensis* | SRR14135877 | 4 | Transcriptome |
| Pakistanapseudes_sp | Tanaidacea | *Pakistanapseudes* sp. | SRR14135870 | 5 | Transcriptome |
| S_cokei | Mysida | *Stygiomysis cokei* | SRR5140123 | 4 | Transcriptome |
| Apseudomorpha_sp | Tanaidacea | | SRR14135860 | 3 | Transcriptome |
| Leptochelia_sp | Tanaidacea | *Leptochelia* sp. | SRR5140116 | 3 | Transcriptome |
| P_assaforensis | Isopoda | *Proasellus assaforensis* | ERR1437543 | 2 | Transcriptome |
| P_jaloniacus | Isopoda | *Proasellus jaloniacus* | ERR1433114 | 3 | Transcriptome |

| | | | | | |
|---|---|---|---|---|---|
| P_margalefi | Isopoda | *Proasellus margalefi* | ERR1437537 | 3 | Transcriptome |
| P_solanasi | Isopoda | *Proasellus spolanasi* | ERR1437540 | 3 | Transcriptome |
| A_alascensis | Tanaidacea | *Arctotanais alascensis* | SRR14135869 | 4 | Transcriptome |
| N_kuroshio | Tanaidacea | *Neotanais* cf. *kuroshio* | SRR14135871 | 2 | Transcriptome |
| P_cantabricus | Isopoda | *Proasellus cantabricus* | ERR1437554 | 2 | Transcriptome |
| P_ebrensis | Isopoda | *Proasellus ebrensis* | ERR1437553 | 2 | Transcriptome |
| P_littoralis | Tanaidacea | *Paradoxapseudes littoralis* | SRR14135865 | 2 | Transcriptome |
| T_kommritzia | Tanaidacea | *Tanaella kommritzia* | SRR14135867 | 3 | Transcriptome |
| A_bangokensis | Bathynellacea | *Allobathynella bangokensis* | SRR4198915 | 0 | Transcriptome |
| Akanthophoreus_sp | Tanaidacea | *Akanthophoreus* sp. | SRR14135881 | 1 | Transcriptome |
| C_sinusa | Tanaidacea | *Chauliopleona* cf. *sinusa* | SRR14135868 | 1 | Transcriptome |
| D_hoyi | Amphipoda | *Diporeia hoyi* | SRR5341787 | 2 | Transcriptome |
| Heterotanoides_sp | Tanaidacea | *Heterotanoides* sp. | SRR14135866 | 1 | Transcriptome |
| Siphonolabrum_sp | Tanaidacea | *Siphonolabrum* sp. | SRR14135862 | 1 | Transcriptome |
| Tanaopsis_sp | Tanaidacea | *Tanaopsis* sp. | SRR14135863 | 1 | Transcriptome |
| B_molinai | Isopoda | *Bragasellus molinai* | ERR1437534 | 6 | Transcriptome |
| Chondrochelia_sp | Tanaidacea | *Chondrochelia* sp. | SRR14135861 | 1 | Transcriptome |
| H_sasuke | Tanaidacea | *Zeuxo ezoensis* | SRR14135878 | 0 | Transcriptome |
| N_indica | Isopoda | *Norileca indica* | SRR18883214 | 0 | Transcriptome |
| Parakanthophoreus_sp | Tanaidacea | *Parakanthophoreus* sp. | SRR14135864 | 0 | Transcriptome |
| Paranarthrura_sp | Tanaidacea | *Paranarthrura* sp. | SRR14135859 | 1 | Transcriptome |
| Sinelobus_sp | Tanaidacea | *Sinelobus* sp. | SRR14135875 | 0 | Transcriptome |
| Z_ezoensis | Tanaidacea | *Zeuxo ezoensis* | SRR14135882 | 1 | Transcriptome |
| Acanthonotozomoides_oatesi_Aoat_MCZ172843 | Amphipoda | *Acanthonotozomoides oatesi* | MCZ172843 | 477 | UCE Library |
| Eusiridae_Eusi1_MCZ172955 | Amphipoda | | MCZ172955 | 465 | UCE Library |
| Podocerus_brasiliensis_PbraSI_USNM1605825 | Amphipoda | *Podocerus brasiliensis* | USNM1605825 | 224 | UCE Library |
| Cyphocaris_richardi_Crich_MCZ172829 | Amphipoda | *Cyphocaris richardi* | MCZ172829 | 484 | UCE Library |

| | | | | | |
|---|---|---|---|---|---|
| Deutella_incerta_Dinc_USNM1605057 | Amphipoda | *Deutella incerta* | USNM1605057 | 254 | UCE Library |
| Hyachelia_tortugae_Htor1_MCZ173266 | Amphipoda | *Hyachelia tortugae* | MCZ173266 | 380 | UCE Library |
| Cyamus_boopis_CybBW1_MCZ173261 | Amphipoda | *Cyamus boopis* | MCZ173261 | 251 | UCE Library |
| Neoxenodice_sp_Neox_MCZ172934 | Amphipoda | *Neoxenodice* sp. | MCZ172934 | 274 | UCE Library |
| Idotea_sp_Idot_MCZ166902 | Isopoda | *Idotea* sp. | MCZ166902 | 130 | UCE Library |
| Diastylidae_sp1_Dias1_MCZ172889 | Cumacea | *Diastylidae* sp. 1 | MCZ172889 | 118 | UCE Library |
| Parandania_boecki_Pboe_MCZ172673 | Amphipoda | *Parandania boecki* | MCZ172673 | 345 | UCE Library |
| Serolidae_Sero2_MCZ172956 | Isopoda | | MCZ172956 | 129 | UCE Library |
| Natatolana_sp_Nata_MCZ172842 | Isopoda | *Natatolana* sp. | MCZ172842 | 124 | UCE Library |
| Haploniscus_sp_Haplo_MCZ172849 | Isopoda | *Haploniscus* sp. | MCZ172849 | 98 | UCE Library |
| Eurycope_sp1_Eury1_MCZ172910 | Isopoda | *Eurycope* sp. | MCZ172910 | 105 | UCE Library |
| Munnopsidae_sp2_Munno2_MCZ172914 | Isopoda | *Munnopsidae* sp. | MCZ172914 | 108 | UCE Library |
| Vibilia_sp_Vibi_MCZ172966 | Amphipoda | *Vibilia* sp. | MCZ172966 | 130 | UCE Library |
| Acanthaspidiidae_Actp_MCZ172845 | Isopoda | | MCZ172845 | 104 | UCE Library |
| Vanhoeffenura_sp_Vanh_MCZ172827 | Isopoda | *Vanhoeffenura* sp. | MCZ172827 | 95 | UCE Library |
| Lampropidae_Lamp1_MCZ172887 | Cumacea | | MCZ172887 | 76 | UCE Library |
| Serolidae_sp1_Sero1_MCZ172911 | Isopoda | | MCZ172911 | 114 | UCE Library |
| Munnidae_sp1_Munni1_MCZ172905 | Isopoda | | MCZ172905 | 99 | UCE Library |
| Desmosomatidae_Desmo1_MCZ172908 | Isopoda | | MCZ172908 | 106 | UCE Library |
| Mesosignidae_Meso1_MCZ172923 | Isopoda | | MCZ172923 | 86 | UCE Library |
| Gnathiidae_Gnath_MCZ172918 | Isopoda | | MCZ172918 | 80 | UCE Library |
| Dyopedos_sp_Dyop_USNM1644295 | Amphipoda | *Dyopedos* sp. | USNM1644295 | 135 | UCE Library |
| Rocinela_sp_Roci_MCZ173268 | Isopoda | *Rocinela* sp. | MCZ173268 | 86 | UCE Library |
| Themisto_gaudichaudii_Tgau_MCZ172670 | Amphipoda | *Themisto gaudichaudii* | MCZ172670 | 113 | UCE Library |
| Tanaidacea_TanANT_MCZ17292 | Tanaidacea | | MCZ17292 | 91 | UCE Library |
| Mictocaris_halope_Mhal_MCZ135174 | Mictacea | *Mictocaris halope* | MCZ135174 | 84 | UCE Library |
| Ischnomesidae_sp2_Ischno2_MCZ172930 | Isopoda | | MCZ172930 | 89 | UCE Library |
| Zeuxo_sp_Zeux1_MCZ173265 | Tanaidacea | *Zeuxo* sp. | MCZ173265 | 63 | UCE Library |
| Laticorophium_baconi_Lbac_USNM1605092 | Amphipoda | *Laticorophium baconi* | USNM1605092 | 148 | UCE Library |
| Nebalia_longicornis_Nlon_MCZ172977 | Leptostraca | *Nebalia longicornis* | MCZ172977 | 91 | UCE Library |
| Antarctomysis_maxima_Amax_MCZ140283 | Mysida | *Antarctomysis maxima* | MCZ140283 | 87 | UCE Library |
| Thermosbaenacea_Thermo_MCZ134138 | Thermosbaenacea | | MCZ134138 | 58 | UCE Library |
| Ischnomesidae_sp1_Ischno1_MCZ172929 | Isopoda | | MCZ172929 | 91 | UCE Library |
| Erythropinae_Eryt_MCZ172927 | Mysida | | MCZ172927 | 97 | UCE Library |
| Caprella_penantis_Cpen_USNM1463333 | Amphipoda | | USNM1463333 | 250 | UCE Library |

| | | | | | |
|---|---|---|---|---|---|
| Epimeria_sp_Epim1_MCZ172924 | Amphipoda | *Epimeria* sp. | MCZ172924 | 183 | UCE Library |
| Nannastacidae_Nanna1_MCZ172885 | Cumacea | | MCZ172885 | 56 | UCE Library |
| Ampeliscidae_Ampe1_MCZ172983 | Amphipoda | | MCZ172983 | 141 | UCE Library |
| Hyperia_curvicephala_Hypcur_MCZ172965 | Amphipoda | *Hyperia curvicephala* | MCZ172965 | 75 | UCE Library |
| Glyptonotus_antarcticus_Gant_MCZ140281 | Isopoda | *Glyptonotus antarcticus* | MCZ140281 | 73 | UCE Library |
| Ianthopsis_sp_Janto_MCZ172931 | Isopoda | *Ianthopsis* sp. | MCZ172931 | 35 | UCE Library |
| Gnathophausia_zoea_Gzoe_MCZ46341 | Lophogastrida | *Gnathophausia zoea* | MCZ46341 | 43 | UCE Library |
| Phronima_sp_Phro_MCZ173260 | Amphipoda | *Phronima* sp. | MCZ173260 | 50 | UCE Library |
| Eurythenes_obesus_Eobes_MCZ172669 | Amphipoda | *Eurythenes obesus* | MCZ172669 | 28 | UCE Library |
| Aporobopyrina_anomala_Aano_MCZ80803 | Isopoda | *Aporobopyrina anomala* | Aano_MCZ80803 | 12 | UCE Library |
| Eucopia_unguiculata_Eung_MCZ37895 | Lophogastrida | *Eucopia unguiculata* | MCZ37895 | 2 | UCE Library |
| Excorallana_sp_Exc_MCZ173267 | Isopoda | *Excorallana* sp. | MCZ173267 | 10 | UCE Library |
| Speleomysis_quinterensis_Squi_MCZ38530 | Stygiomysida | *Speleomysis quinterensis* | MCZ38530 | 4 | UCE Library |
| Diastylis_polita_Dipo1_MCZ35770 | Cumacea | *Diastylis polita* | MCZ35770 | 6 | UCE Library |
| Tritella_laevis_Tlae_USNM1581261 | Amphipoda | *Tritella laevis* | USNM1581261 | 0 | UCE Library |
| Anaspides_tasmaniae_Atas_MCZ10400 | Anaspidacea | *Anaspides tasmaniae* | MCZ10400 | 0 | UCE Library |
| Apseudes_neotanais_Aneo_MCZ48367 | Tanaidacea | *Apseudes neotanais* | MCZ48367 | 3 | UCE Library |
| Caprella_horrida_Chor1_MCZ173197 | Amphipoda | *Caprella horrida* | MCZ173197 | 3 | UCE Library |
| Dulichia_tuberculata_Dtur1_MCZ173223 | Amphipoda | *Dulichia tuberculata* | MCZ173223 | 2 | UCE Library |
| Eudorella_truncatula_Etru_MCZ57214 | Cumacea | *Eudorella truncatula* | MCZ57214 | 1 | UCE Library |
| Neomysis_americana_Name_MCZ44388 | Mysida | *Neomysis americana* | MCZ44388 | 0 | UCE Library |
| Potiicoara_brasiliensis_Poti1_MCZ173273 | Spelaeogriphacea | *Potiicoara brasiliensis* | MCZ173273 | 1 | UCE Library |
| Apseudes_nosphyrapus_Anos_MCZ48369 | Tanaidacea | | MCZ48369 | 2 | UCE Library |
| Cercops_sp_Cerc1_MCZ173239 | Amphipoda | *Cercops* sp. | MCZ173239 | 1 | UCE Library |
| Chondrochelia_brasiliensis_Chon1__MCZ173264 | Tanaidacea | *Chondrochelia brasiliensis* | MCZ173264 | 4 | UCE Library |
| Gnathophausia_gracilis_Ggra1_MCZ173276 | Lophogastrida | *Gnathophausia gracilis* | MCZ173276 | 1 | UCE Library |
| Gnathophausia_ingens_Ging1_MCZ173274 | Lophogastrida | *Gnathophausia ingens* | MCZ173274 | 1 | UCE Library |
| Lophogaster_sp_Lopho2_MCZ173275 | Lophogastrida | *Lophogaster* sp. | MCZ173275 | 1 | UCE Library |
| Caprella_mendax_Cmen_USNM1581205 | Amphipoda | *Caprella mendax* | USNM1581205 | 1 | UCE Library |
| Dulichia_wolffi_Dwol1_MCZ173224 | Amphipoda | *Dulichia wolffi* | MCZ173224 | 1 | UCE Library |
| Dyopedos_knipowitschi_Dkni1_MCZ173220 | Amphipoda | *Dyopedos knipowitschi* | MCZ173220 | 0 | UCE Library |
| Dyopedos_monacanthus_Dmon1_MCZ173216 | Amphipoda | *Dyopedos monacanthus* | MCZ173216 | 0 | UCE Library |

| | | | | | |
|---|---|---|---|---|---|
| Dyopedos_normani_Dnor1_MCZ173217 | Amphipoda | *Dyopedos normani* | MCZ173217 | 0 | UCE Library |
| Dyopedos_porrectus_Dpor_MCZ173225 | Amphipoda | *Dyopedos porrectus* | MCZ173225 | 0 | UCE Library |
| Dyopedos_spinosus_Dspi1_MCZ173218 | Amphipoda | *Dyopedos spinosus* | MCZ173218 | 0 | UCE Library |
| Ingolfiellida_Ingol2_MCZ79594 | Ingolfiellida | | MCZ79594 | 0 | UCE Library |
| Paradulichia_typica_Ptyp1_MCZ173226 | Amphipoda | *Paradulichia typica* | MCZ173226 | 0 | UCE Library |

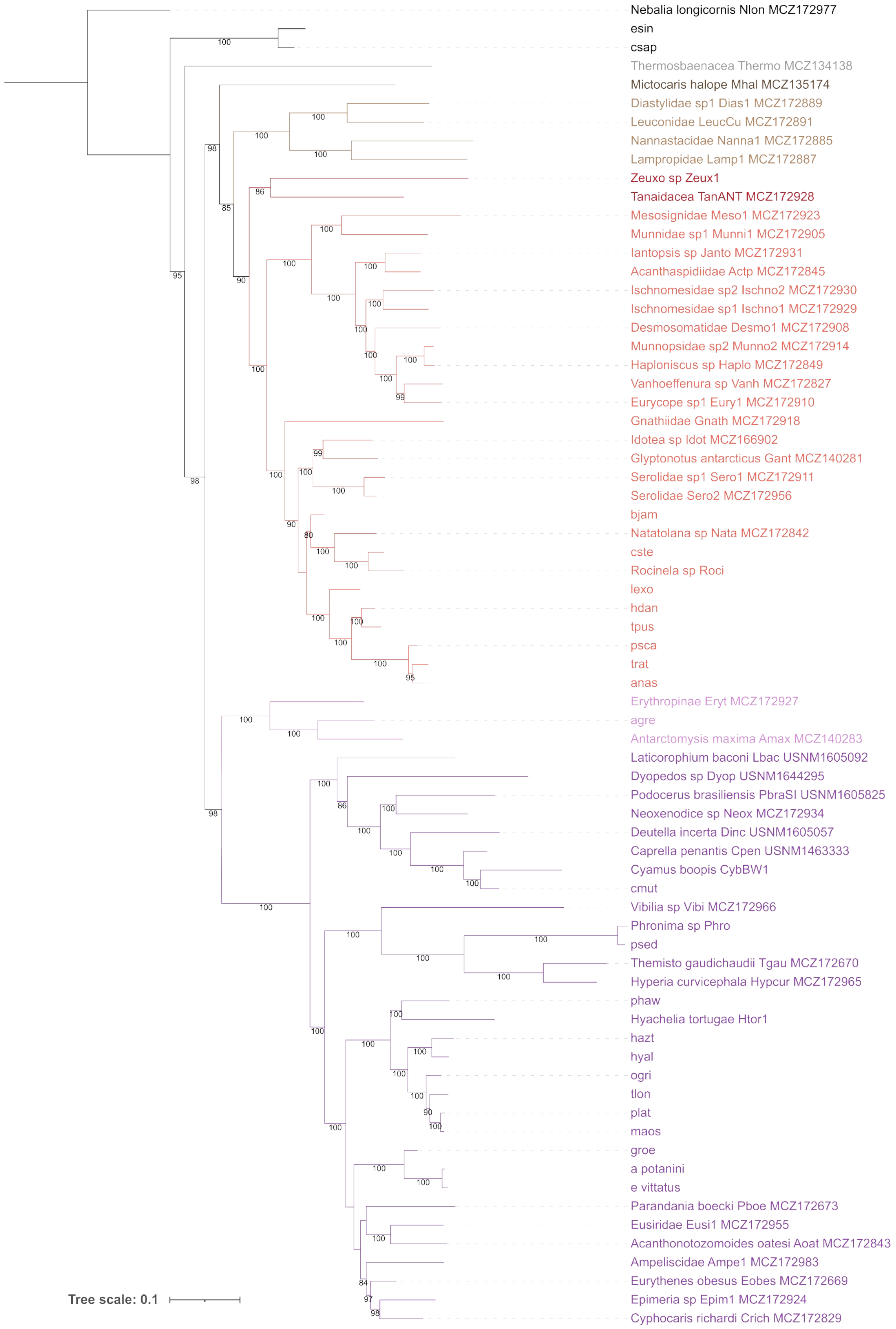


**Figure S1.** Maximum likelihood tree inferred from partitioned analysis of concatenated M35 dataset in IQTREE.

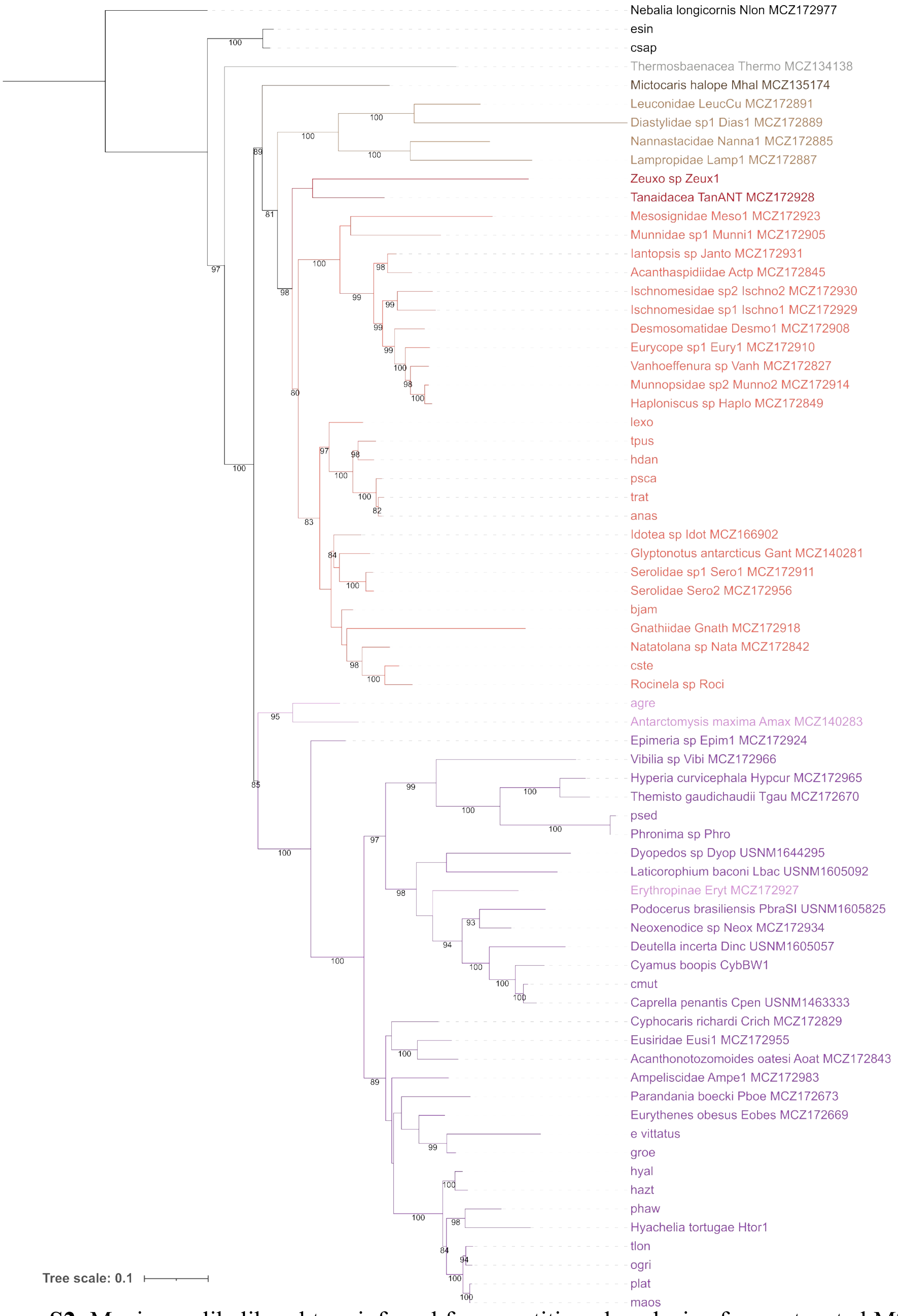


**Figure S2.** Maximum likelihood tree inferred from partitioned analysis of concatenated M75 dataset in IQTREE.

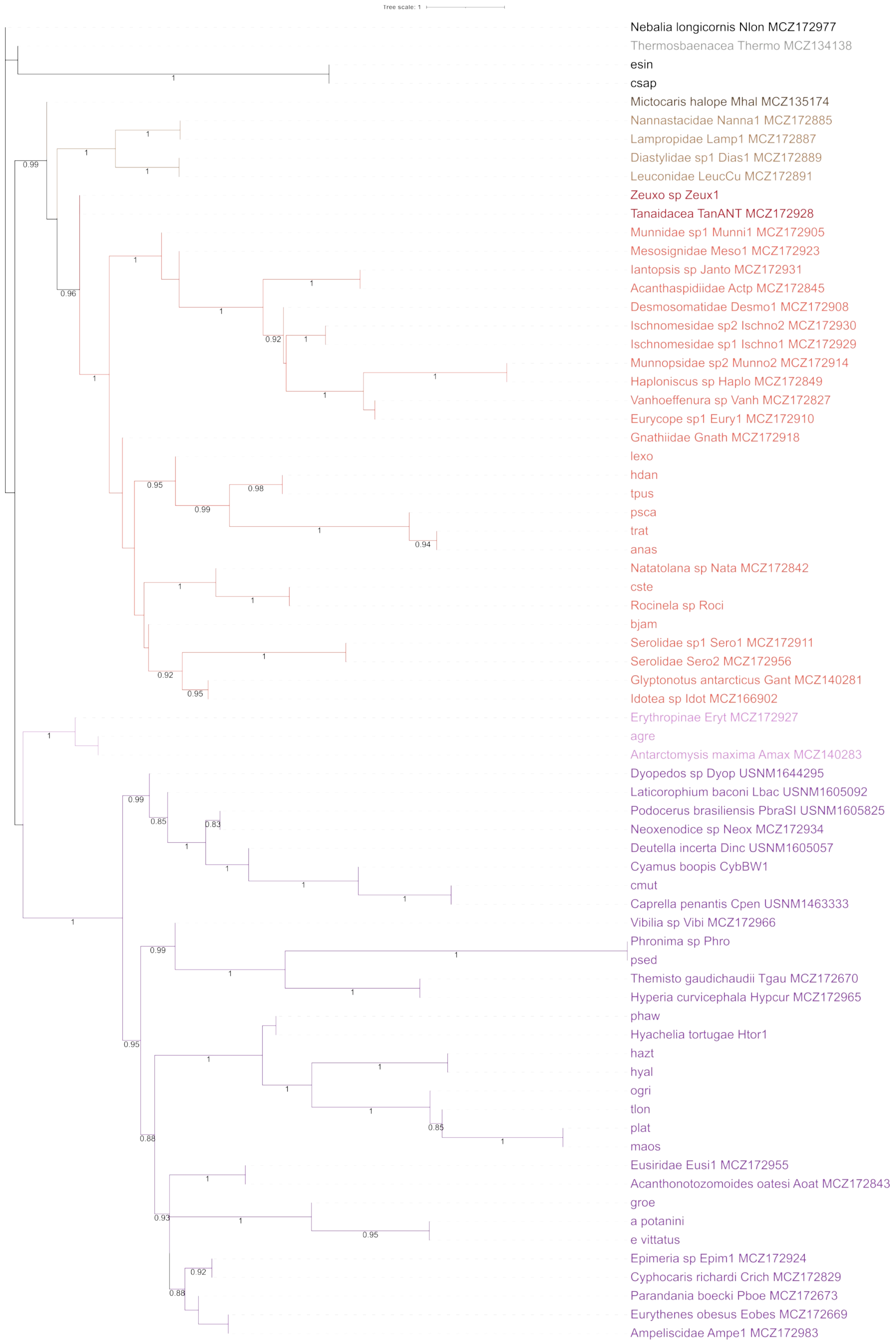


**Figure S3.** Coalescent-based tree inferred from M35 dataset in ASTRAL.

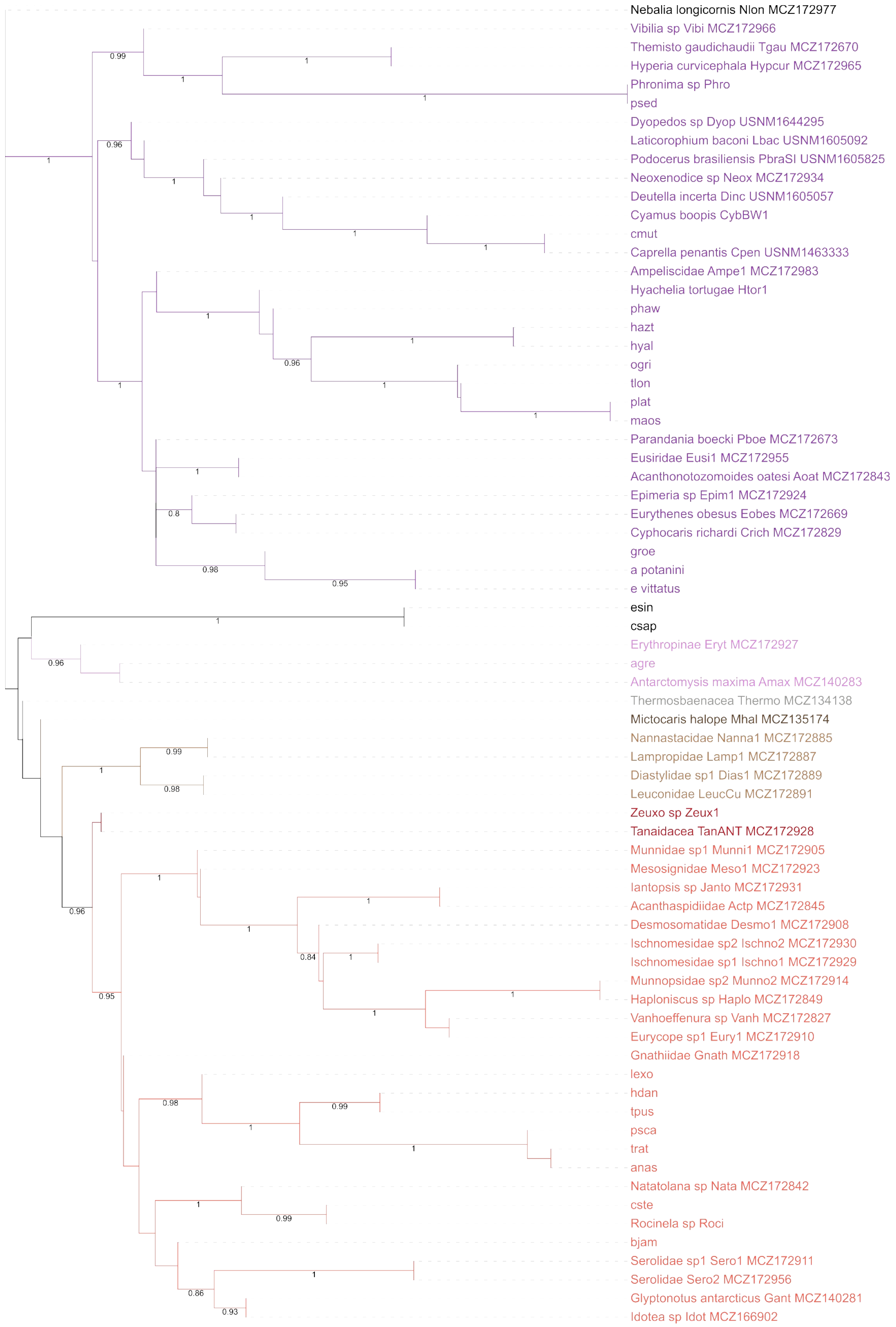


**Figure S4.** Coalescent-based tree inferred from M50 dataset in ASTRAL.

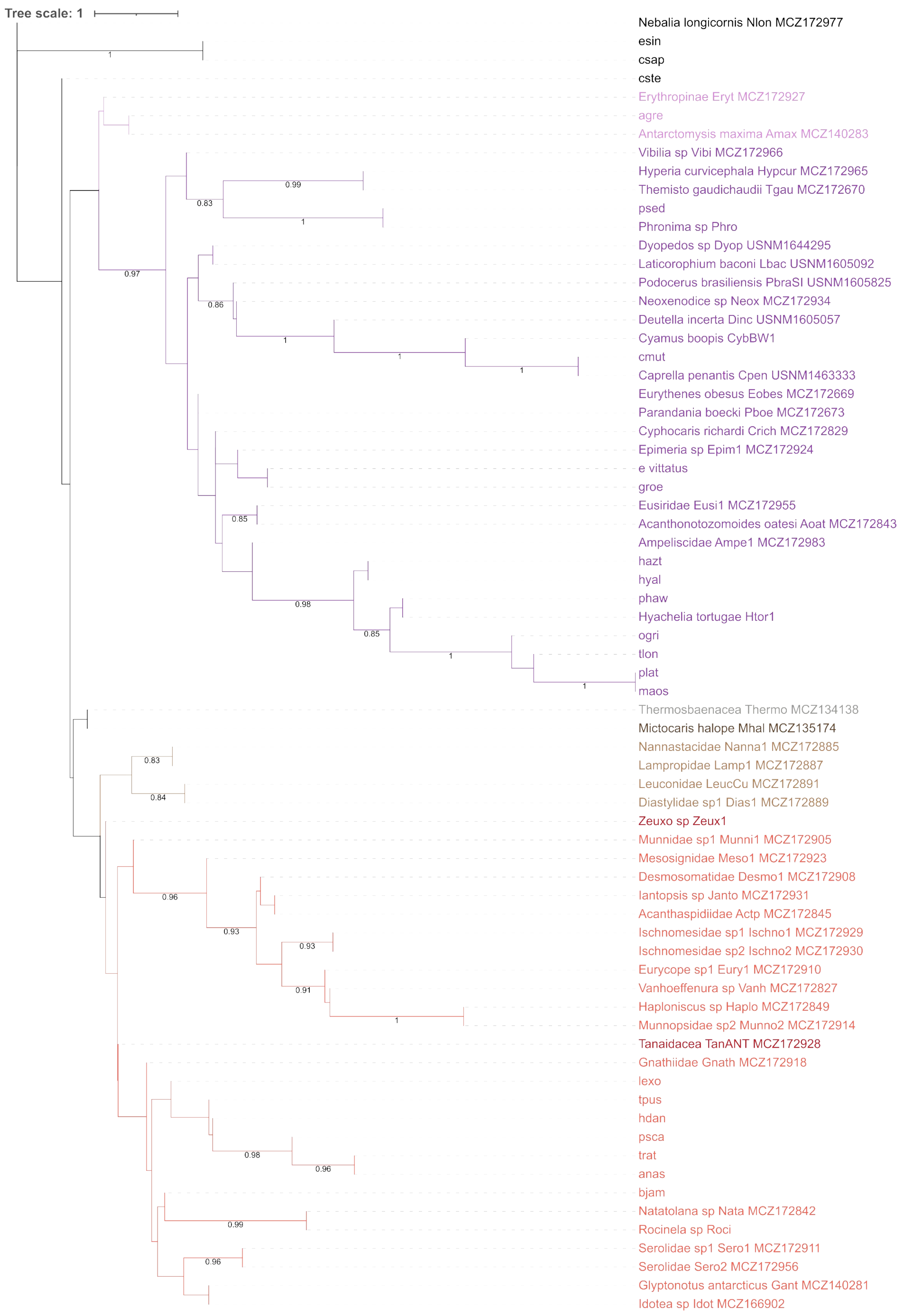


**Figure S5.** Coalescent-based tree inferred from M75 dataset in ASTRAL.
